# Root-associated Fungi in Orchidaceae: Diversity, Phylogeny, Ecology, and Outstanding Questions

**DOI:** 10.1101/2022.12.16.519622

**Authors:** Deyi Wang, Jun Lerou, Jorinde Nuytinck, Sofia I.F. Gomes, Hans Jacquemyn, Vincent S.F.T. Merckx

**Affiliations:** Naturalis Biodiversity Center, Leiden, the Netherlands; Institute of Biology, Leiden University, Leiden, the Netherlands; Department of Biology, Plant Conservation and Population Biology, KU Leuven, Leuven, Belgium; Department of Evolutionary and Population Biology, Institute for Biodiversity and Ecosystem Dynamics, University of Amsterdam, Amsterdam, the Netherlands

**Keywords:** Orchid mycorrhiza, Phylogeny, *Basidiomycota*, *Ascomycota*, Fungal lifestyle, Orchidaceae

## Abstract

Mycorrhizal fungi form ubiquitous symbiotic associations with almost all land plants and are of key interest to evolutionary biologists and ecologists because this ancient symbiosis was essential for the colonization of land by plants – a major turning point in the evolutionary history of the earth – and the subsequent development and functioning of the terrestrial ecosystems. Within the orchid family (Orchidaceae), plants establish unique interactions with specific orchid mycorrhizal fungi. These fungal symbionts are essential for the development of orchids as they provide carbon and soil nutrients to germinating orchid seeds and the nutritional supply continues for adult orchids to different degrees. Fueled by the development of DNA sequencing techniques, the diversity of mycorrhizal and other root-associated fungi in orchid roots has been extensively reported in evolutionary and ecophysiological studies. However, the full taxonomic range of orchid-associated fungi remains to be investigated in a broad phylogenetic framework, hampering a further understanding of the evolution and ecological adaptation of orchid mycorrhizal interactions. In this study, we used the most complete DNA dataset to date to map the phylogenetic distribution and ecological lifestyles of root-associated fungi in Orchidaceae by phylogenetic reconstructions at the fungal order level. We found that a broad taxonomic range of fungi (clustered into 1898 operational taxonomic units) resided in orchid roots, belonging to at least 150 families in 28 orders in *Basidiomycota* and *Ascomycota*. These fungi were assigned to diverse ecological lifestyles including typical orchid mycorrhizal fungi (‘rhizoctonia’), ectomycorrhizal fungi, wood- or litter-decaying saprotrophic fungi, and other endophytes/pathogens/saprotrophs. This overview reveals that among the four different mycorrhizal types, the orchid mycorrhizal symbiosis probably involves the highest diversity of fungal taxa. We hope that our newly reconstructed phylogenetic framework of orchid-associated fungi and the assessment of their potential mycorrhizal status will benefit future ecological and evolutionary studies on orchid-fungal interactions.

## 1 Introduction

Mycorrhizal symbiosis is an ancient and ubiquitous association between plants and root- or rhizoid-associated mycorrhizal fungi (Frank & Trappe 2005; Smith & Read 2008). In this generally mutualistic symbiosis, mycorrhizal fungi promote plant growth and fitness by facilitating the acquisition of water and mineral nutrients from the soil and by providing protection against abiotic stresses and pathogens. In return, the plant hosts provide carbon derived from photosynthesis to the fungal symbionts (Smith & Read 2008). Mycorrhizal fungi are widespread and can be found in almost all ecosystems from forests to deserts and arable lands (Read 1991; Brundrett 2009; Ji & Bever 2012). It has been estimated that over 90 % of all extant land plant species establish associations with mycorrhizal fungi (Wang & Qiu 2006; van der Heijden et al. 2015; Brundrett & Tedersoo 2018). Based on the morphology of the interaction and the identity of the partners, four major mycorrhizal types have been described: arbuscular mycorrhizas, ectomycorrhizas, ericoid mycorrhizas, and orchid mycorrhizas (Smith & Read 2008; van der Heijden et al. 2015; Tedersoo et al. 2020). Arbuscular mycorrhizal fungi belong to *Glomeromycotina* and *Mucoromycotina* (*Mucoromycota*) (Öpik et al. 2010; Bidartondo et al. 2011; Hoysted et al. 2018), ectomycorrhizal fungi include ca. 20,000 species in ~250 genera in *Basidiomycota* and *Ascomycota* (Tedersoo et al. 2010; Tedersoo & Smith 2013, 2017; van der Heijden et al. 2015) or an estimation of 20,000 – 25,000 species based on knowns and unknows in macromycete diversity (Rinaldi et al. 2008; Comandini et al. 2012), and ericoid mycorrhizal fungi represent a small group of fungi in *Leotiomycetes (Ascomycota*) (Walker et al. 2011; Kohout 2017). With an estimated 25,000 taxa across *Basidiomycota* and *Ascomycota*, orchid mycorrhizal fungi rival the diversity of ectomycorrhizal fungal taxa (van der Heijden et al. 2015). The estimated vast diversity of orchid mycorrhizal fungi has been attributed to the high specificity of orchid mycorrhizal associations (Dearnaley et al. 2012; Martos et al. 2012; van der Heijden et al. 2015) and the extraordinarily high plant diversity (>27,000 species) involved in the symbiosis, as well as the specialized niche of orchids across the globe (Chomicki et al. 2015; Givnish et al. 2015).

Orchid mycorrhizal fungi are critical for the development and growth of orchids (Dressler & Rasmussen 1996; Taylor et al. 2002; Smith & Read 2008). Under natural conditions, mycorrhizal fungi provide essential carbon (C) and mineral nutrients to the endosperm-lacking seeds of orchids for germination, and for their underground protocorms and tubers to grow into adults in a phenomenon called initial mycoheterotrophy (Merckx 2013; Jacquemyn & Merckx 2019). Most orchids become autotrophic after they appear above the ground and trade photosynthetically derived C for organic and inorganic nutrients with their fungal partners (Cameron et al. 2006, 2008). In contrast to autotrophic adult orchids, some orchids continue to depend fully or partially on fungal C. These orchid species are termed full and partial mycoheterotrophs, respectively (Merckx 2013). Despite the traditional view that mycoheterotrophic orchids do not supply carbon to their mycorrhizal fungi, they may provide other benefits (ammonium, vitamins, and biotic/abiotic protections) to fungal partners to maintain a symbiotic mutualism (Dearnaley et al. 2012; Dearnaley & Cameron 2017; Fochi et al. 2017; Yeh et al. 2019).

The detection and identification of fungal symbionts in orchid roots rely on various morphological, physiological, and molecular methods. Orchid mycorrhizal fungi form elaborate coiled hyphae known as pelotons in the root cortex cells of orchids (Smith & Read 2008; Dearnaley et al. 2012). The microscopic observation of fungal pelotons within orchid root cells is the most traditional and reliable way to confirm the establishment of fungal colonization. Symbiotic seed germination experiments and fungal cultures have long been adopted to confirm the mycorrhizal status of the isolated fungi (Warcup 1971; Clements et al. 1986; Xu & Guo 1989; Rasmussen 1995). However, the development of molecular techniques in the late twentieth century has facilitated the detection and identification of fungi that are difficult to culture or hard to identify based on morphological characteristics (Bruns et al. 1998; Horton & Bruns 2001; Dearnaley 2007). With a combination of morphological, physiological, and molecular methods on orchid mycorrhizal fungi, it has become clear that most orchids associate with so-called ‘rhizoctonia’ fungi, a polyphyletic group including three distinct families in the phylum *Basidiomycota*: *Tulasnellaceae* and *Ceratobasidiaceae* (in the *Cantharellales*), and *Serendipitaceae* (in the *Sebacinales*) (Smith & Read 2008; Yukawa et al. 2009; Dearnaley et al. 2012; Rasmussen et al. 2015). These fungal taxa are generally soil-dwelling saprobes except that a few subclades form orchid mycorrhizas, as well as ectomycorrhizas (Rinaldi et al. 2008; Tedersoo et al. 2010; Tedersoo & Smith 2013, 2017) and ericoid mycorrhizas (van der Heijden et al. 2015). Rhizoctonias are thought to have established a symbiotic association with the common ancestor of all extant orchids (Yukawa et al. 2009; Wang et al. 2021; Selosse et al. 2022), representing a major evolutionary innovation from ancestral arbuscular associations (van der Heijden et al. 2015; Brundrett & Tedersoo 2018; Feijen et al. 2018).

Although rhizoctonia fungi are prevalent orchid mycorrhizal fungi, many orchids also associate with non-rhizoctonia fungi of *Basidiomycota* and *Ascomycota*. This wide range of non-rhizoctonia fungal taxa has mainly been detected with molecular techniques, including recent high-throughput sequencing platforms (Dearnaley et al. 2012; Jacquemyn et al. 2016, 2017, 2021). Most of these taxa have a saprotrophic or ectomycorrhizal lifestyle and are assumed to be secondarily recruited by orchids as mycorrhizal partners (Selosse et al. 2022). Ectomycorrhizal fungi (mostly reported in *Russulaceae*, *Sebacinaceae*, and *Thelephoraceae*) simultaneously link to surrounding plants within common mycelium networks with orchids, in which the C resources from the surrounding vegetation are ultimately transported to orchids (Dearnaley et al. 2012; Merckx 2013). The C transfer through fungal networks to orchids has been confirmed by isotope tracing, fungal inoculation, and pot culture experiments (Zelmer & Currah 1995; McKendrick et al. 2000a; Bougoure et al. 2010; Yagame & Yamato 2013). Major clades of wood- and litter-decaying saprotrophic fungi have been detected in orchid roots, mostly within fully mycoheterotrophic orchids, including members of *Agaricales*, *Hymenochaetales*, and *Polyporales* (Dearnaley et al. 2012; Lee et al. 2015; Ogura-Tsujita et al. 2018, 2021). These saprotrophic fungi feature higher activities of lignocellulose-degrading enzymes than ectomycorrhizal fungi (Kohler et al. 2015) and exhibit distinctive isotope signatures (Lee et al. 2015; Schiebold et al. 2017; Ogura-Tsujita et al. 2018).

In addition, some fungal taxa that have been frequently or occasionally detected in orchid roots by molecular sequencing have been largely discounted as ‘molecular scraps’ (Bayman & Otero 2007; Selosse et al. 2010) and filtered out according to the taxonomic range of the rhizoctonia complex and/or a list of putative orchid mycorrhizal fungi (Dearnaley et al. 2012). For example, members of *Helotiales, Pezizales, Xylariales, Pleosporales*, and *Hypocreales* that have been frequently detected in orchid roots were assumed to be non-mycorrhizal with orchids (Bayman & Otero 2007; Tao et al. 2008; Oliveira et al. 2014). Yet, their non-mycorrhizal status in orchids deserves further confirmation since at least some of these fungi resemble typical orchid mycorrhizal fungi by forming peloton-like hyphal structures within root cells (Hou & Guo 2009; Jiang et al. 2019; Sisti et al. 2019) or by promoting seed germination and seedling development of several orchids (Vujanovic 2000; Ma et al. 2015). Better insights into the phylogenetic range of these endophytes may allow tracing potential evolutionary transitions to novel mycorrhizal associations with orchids (Selosse et al. 2022).

Over the past decade, a plethora of data on fungal taxa that form symbiotic associations with orchid roots has been generated, but these data have yet to be integrated within a single phylogenetic framework. The major aim of this study is to provide a broad overview of the phylogenetic distribution and ecological functions of orchid root-associated fungi. To facilitate readability, we use the general term ‘OrM fungi’ to include both orchid mycorrhizal fungi and putatively non-mycorrhizal fungi dwelling in orchid roots. Using a newly compiled dataset of fungi detected in orchid roots based on eco-physiological studies and on molecular sequencing of orchid mycorrhizas (Wang et al. 2021), we explore the potential taxonomic range and ecological lifestyles of OrM fungi of the Orchidaceae. First, we investigate the taxonomic range of OrM fungi by phylogenetic reconstructions within each fungal order. Furthermore, we examine their ecological lifestyles, including ‘rhizoctonias’, ectomycorrhizal fungi, non-rhizoctonia saprotrophic fungi, and other endophytic/pathogenic/saprotrophic lifestyles. Finally, based on our assessment on taxonomic diversity and ecological roles of orchid-associated fungi, we provide implications for future studies on the evolution and ecophysiology of orchid-fungal interactions in general.

## 2 Materials and Methods

### 2.1 Molecular dataset, sequence filtering, and taxonomic assignment

Fungal ITS accessions were extracted from a newly built molecular dataset of orchid mycorrhiza (Table S1 in Wang et al. (2021)). This dataset contains 8860 accessions of fungal internal transcribed spacer (ITS) region detected in roots of 750 orchid species covering all five subfamilies of Orchidaceae, which were sampled from 50 countries and regions across the globe. The top 10 countries/regions had over 80% of sequences, including USA (16.3%), Belgium (13.7%), Italy (10.7%), China (8.7%), Australia (7.5%), Reunion (6.1%), Ecuador (5.7%), Japan (5.4%), Brazil (4.1%), and UK (3.5%). ITS sequences recorded in this dataset were generated by both traditional sequencing approaches (77.8% of sequences by Sanger sequencing, and DNA arrays) and high-throughput sequencing platforms (22.2% of sequences by 454 amplicon pyrosequencing and Illumina Miseq platforms). For Sanger sequencing and DNA arrays, forward primers ITS1 (White et al. 1990), ITS1F (Gardes & Bruns 1993), or ITS1OF (Taylor & McCormick, 2008) were commonly used in combination with reverse primer ITS4 (White et al. 1990), ITS4OF (Taylor & McCormick, 2008), ITS4Tul (Taylor & McCormick 2008), TW13 (Selosse et al. 2007), or cNL2F (White et al. 1990), generating PCR products spanning both ITS1 and ITS2 regions (generally over 500 bases). For high-throughput sequencing, forward primers ITS86 (Turenne et al., 1999) and ITS3 (Taylor & McCormick, 2008) were usually used for PCR amplification in combination with one of the reverse primers ITS4, ITS4OF, and ITS4Tul, generating PCR products of ca. 300 bases on average and covering the ITS2 region and the adjacent partial 5.8S and 28S rRNAs.

DNA sequences of these accessions were downloaded from the NCBI GenBank database (Sayers et al. 2022) and imported into the Geneious Prime v.2019.2 platform (https://www.geneious.com) to annotate gene regions (18S rRNA, ITS1, 5.8S rRNA, ITS2 and 28S rRNA) using local fungal ITS sequences. Sequences that were not annotated with fungal ITS regions were discarded. Ambiguous bases at two ends of sequences were manually trimmed to improve sequence quality. All filtered 8818 sequences were provided in Appendix S1. After filtering and quality control, fungal sequences were split by order according to the taxonomic assignment (Appendix S2) using USEARCH v11 (Edgar 2010). The subsequent order-based analyses were aimed to avoid ambiguity in operational taxonomic unit (OTU) clustering (Blaxter et al. 2005) and to reduce data size for sequence alignment and phylogenetic inference. The ‘usearch_global’ blast command was used for blast searches of each fungal sequence against the latest reference database of UNITE (utax_reference_dataset_02.02.2019.fasta.gz; Retrieved in January 2020) (UNITE Community 2019). The UNITE dataset uses the NCBI Taxonomy classification as a taxonomic backbone, supplemented with modifications from Index Fungorum and MycoBank (Nilsson et al. 2019).

Fungal orders having less than 10 sequences and in total comprising 1.7% of all filtered sequences were not considered for subsequent OTU analyses. Within each of the remaining orders, sequences were clustered into OTUs with USEARCH based on 97 % sequence similarity. The global singletons and doubletons were kept for further analyses to retain as much taxonomic diversity of orchid mycorrhizal fungi as possible in our dataset. Taxonomic information for OTUs was assigned by USEARCH against the UNITE local database and the hit with the highest identity possible (identity > 90) was retrieved. The dataset of all clustered OTUs of orchid-associated fungi and their taxonomic assignment was provided as supplementary files (Appendix S3 and Table S1).

### 2.2 Phylogenetic analyses

We reconstructed the phylogenetic relationships of the filtered fungal OTUs in orders belonging to *Basidiomycota* and *Ascomycota* because these two phyla comprised the majority (99.3 %) of all OTUs. We built order-level phylogenies because the phylogenetic position of orders in *Basidiomycota* and *Ascomycota* is relatively robust whereas the position and delimitation of families are much less stable (Prieto & Wedin 2013; Zhao et al. 2017; Tedersoo et al. 2018; He et al. 2019). We did not include OTUs that were not assigned to the family level (3.6% of all OTUs) for phylogenetic reconstructions because these non-assigned OTUs usually had a short sequence length or exhibited ambiguous taxonomic assignment to the different families. The outgroup taxa were selected for each order according to phylogenetic information provided in the published literature (see details in Table S2). For each family in a fungal order, at least two sequences, if available, were extracted from the UNITE database as taxonomic references.

Subsequently, within each fungal order, the clustered OTUs, reference sequences of each family, and outgroups were aligned with MUSCLE v.3.8.425 (Edgar 2004) with default parameters in the Geneious platform. The alignment was manually trimmed to have an equal length and to mainly comprise the ITS2 region. Short OTUs that were not well aligned or comprised only a few bases of the ITS2 region were removed from the alignment. The final alignment was used to reconstruct the Maximum likelihood (ML) tree for each order using RAxML v.8.2.12 (Stamatakis 2014) in the CIPRES Science Gateway (Miller et al. 2010). The divergence times of the reconstructed phylogeny of each order were estimated using a penalized likelihood approach with treePL v.1.0 (Smith & O’Meara 2012). The crown age of each order was fixed for time calibration, which was extracted from the literature to represent the minimum age of divergence (Table S2). Finally, time-calibrated trees of fungal orders were combined as a single phylogeny using the ‘bind. tree()’ function in the ‘ape’ R package (Paradis & Schliep 2019) supplied with a backbone phylogeny of fungal orders according to available phylogenetic studies (Prieto & Wedin 2013; Kohler et al. 2015; Zhao et al. 2017; Hongsanan et al. 2017; Tedersoo et al. 2018; He et al. 2019; Mao & Wang 2019). Sequence alignment and phylogenetic tree of each fungal order were provided in Appendix S4.

### 2.3 Ecological categories of fungal taxa

We categorize fungal families into several ecological categories (‘lifestyles’ or ‘guilds’) with information available in the literature (Rinaldi et al. 2008; Comandini et al. 2012; Dearnaley et al. 2012; Rasmussen et al. 2015; Tedersoo & Brundrett 2017; Ogura-Tsujita et al. 2021) supplemented by FUNGuild v1.0 (Nguyen et al. 2016) and FungalTraits v1.2 (Põlme et al. 2020). The three families comprising ‘rhizoctonia’ fungi (*Tulasnellaceae*, *Ceratobasidiaceae*, and *Serendipitaceae*) were categorized as ‘rhizoctonia’ (RHI) lifestyle. Apart from their well-known endophytic niche as symbionts of orchid roots, a few subclades of the three rhizoctonia families have multiple ecological niches including free-living soil-dwelling saprotrophs, and potentially forming ectomycorrhiza (Tedersoo & Smith 2013, 2017) and ericoid mycorrhizas (van der Heijden et al. 2015). In this study, we used the concept of rhizoctonia fungi for the three families specifically representing their ability to form orchid mycorrhizas regardless of their multiple ecological niches. Similarly, families mainly comprising ectomycorrhizal fungi were classified as ectomycorrhizal (ECM) while families that mainly contain wood- or litter-decaying saprotrophic fungi were classified as saprotrophic (SAP) although other ecological niches are sometimes recorded for these families as well. Fungal families having both ECM and SAP fungi were classified as the dual ECM/SAP lifestyle. The remaining families, which mainly comprised endophytes, pathogens, parasites, unspecified saprotrophs, or members of unknown ecological guilds, were assigned to ‘others’.

## 3 Results and Discussion

### 3.1 Phylogeny, taxonomic breadth, and lifestyle of OrM fungi

A total of 1898 OTUs were assigned to 150 fungal families and 28 orders in *Basidiomycota* and *Ascomycota* (Fig. 1, Fig. 2, Table 1, and Table S3). If these documented fungal taxa are truly mycorrhizal with orchids, the taxonomic breadth of orchid mycorrhizal fungi is the most speciose among all major mycorrhizal types (Table 1). The observed wide taxonomic range of putative orchid mycorrhizal fungi is much larger than what was summarized in previous studies (ca. 22 families of *Basidiomycota* and *Ascomycota*) (Dearnaley 2007; Dearnaley et al. 2012; Rasmussen et al. 2015), although the families might not be comparable with the past because of family circumscriptions. Also, the function of several orchid-associated fungal lineages remains to be determined (see below). The phylum *Basidiomycota* comprised the majority of OTUs (71.3 % of all OTUs) and included fourteen different orders. Most OTUs were assigned to the order *Cantharellales* (23.6 %), *Agaricales* (15.6 %), *Sebacinales* (12.8 %), *Russulales* (7.8 %), and *Thelephorales* (5.5 %) (Fig. 2 and Table S2). The phylum *Ascomycota* accounted for 29.7 % of all OTUs, most of which belonged to the orders *Helotiales* (5.5 %), *Pezizales* (4.8 %), and *Hypocreales* (4.3 %) (Fig. 2 and Table S2).

**Fig. 1.**
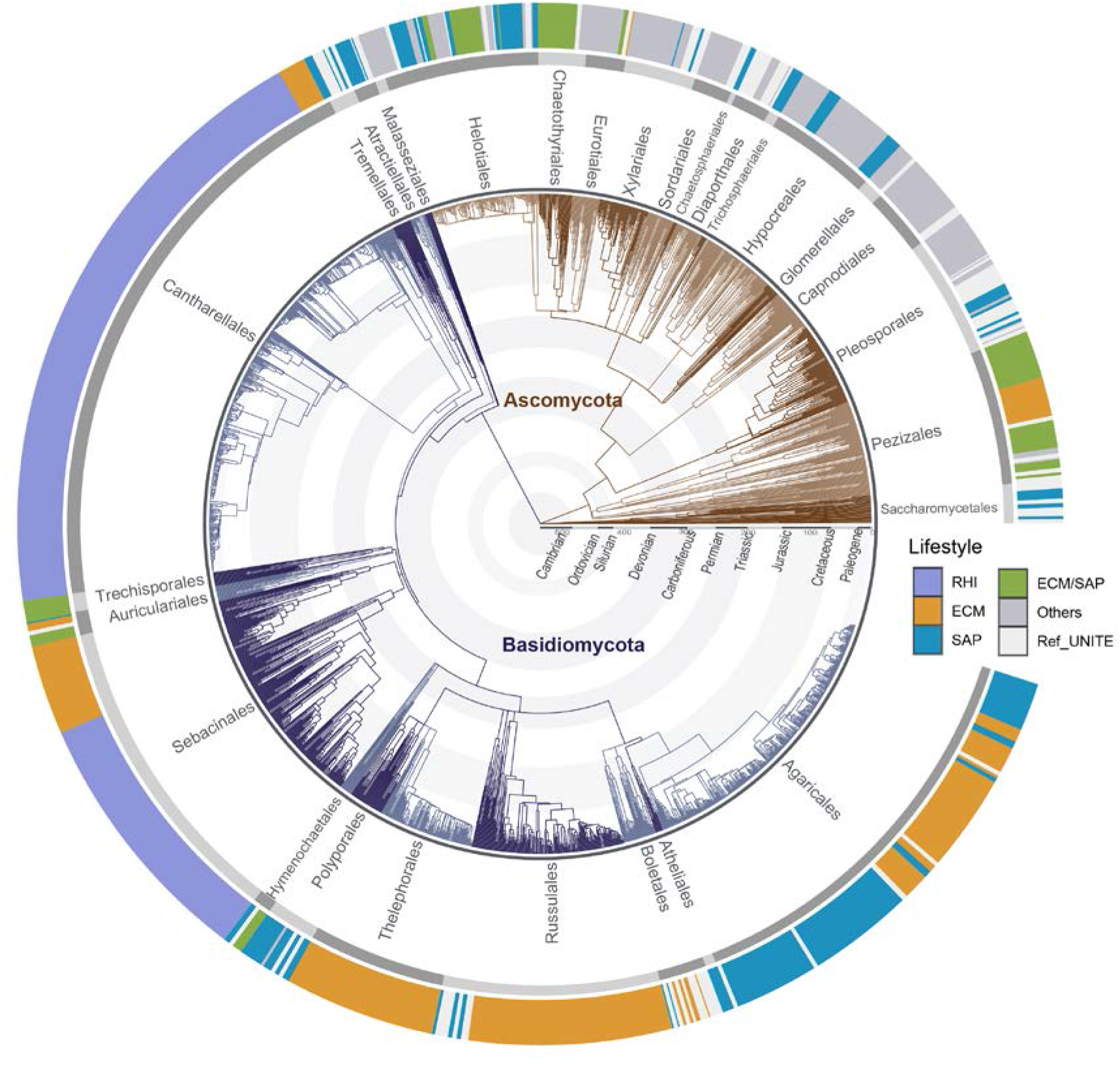
Time-calibrated phylogeny of all fungal OTUs associated with orchid roots and their ecological lifestyles. The fungal phylogeny encompasses all fungal OTUs in *Basidiomycota* and *Ascomycota* based on a newly compiled orchid mycorrhizal dataset (Table S1 in Wang et al. (2021)). The tree branches of *Ascomycota* and *Basidiomycota* are distinguished by brownish and blueish colors. The geological time scales are visualized by circles from the Cambrian to the present. The outer circle annotates the lifestyle of fungal OTUs at the family level. The three fungal families (*Ceratobasidiaceae, Tulasnellaceae*, and *Serendipitaceae*) that comprise typical orchid mycorrhizal fungi were categorized as Rhizoctonia (RHI) fungi. Fungal families mainly comprising ectomycorrhizal or saprotrophic fungi were categorized as the ECM or SAP lifestyle, respectively. Fungal families dominated by both ectomycorrhizal and saprotrophic fungi were classified as ECM/SAP lifestyles. Fungal families mainly comprising endophytes, pathogens, unspecified saprotrophs, or unknown fungi were classified as ‘Others’. The reference sequences from the UNITE database were marked as ‘Ref_UNITE’. To produce a simplified schematic tree for visualization, some tips of reference sequences for phylogenetic reconstructions are not shown in the tree.

**Fig. 2.**
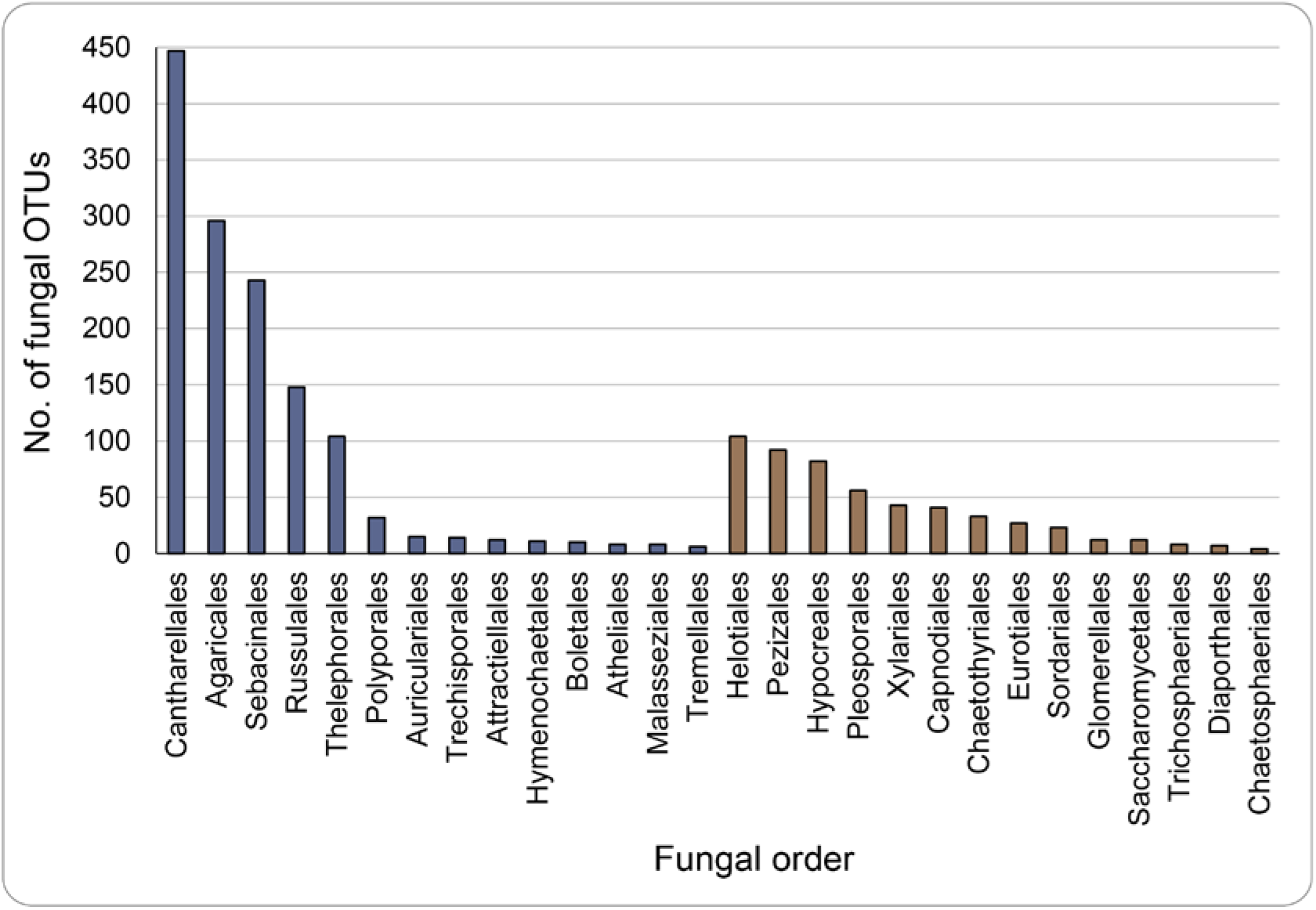
Number of OTUs per order in the phyla *Basidiomycota* (blue) and *Ascomycota* (brown). The fungal orders are ranked by the number of OTUs in each order.

**Table 1.** The taxonomic range of major mycorrhizal fungi.

| Mycorrhizal type | Phylum | Order | Family |
| --- | --- | --- | --- |
| AM | *Mucoromycota*<br>(*Glomeromycotina*) | >6 | >17 |
| ECM | *Basidiomycota* and<br>*Ascomycota* | >18 | >50 |
| ErM | *Basidiomycota* and<br>*Ascomycota* | >4 | >7 |
| OrM | *Basidiomycota* and<br>*Ascomycota* | >28 | >150 |

Note: The numbers of fungal family and orders for AM (arbuscular mycorrhizal fungi), ECM (ectomycorrhizal fungi), and ErM (ericoid mycorrhizal fungi) are according to the FungalTraits v1.2 database (Pölme et al. 2020). The fungal families and orders of OrM fungi are documented based on the orchid mycorrhizal dataset in this study.

Fungal families were annotated with rhizoctonia, ectomycorrhizal, saprotrophic, and other lifestyles (Fig. 3 and Table S3). The three rhizoctonia families comprised the highest number of OTUs (32 % of all OTUs): *Tulasnellaceae* (276 OTUs), *Serendipitaceae* (180 OTUs), and *Ceratobasidiaceae* (149 OTUs). A comparatively large proportion of OTUs (23 % of all OTUs) was assigned to 20 ECM families, followed by 17 % and 7 % of all OTUs assigned to 39 SAP families and 11 ECM/SAP families, respectively. The documented ECM fungi in the orchid mycorrhizal dataset exhibited a wide taxonomic range, accounting for 12/18 ECM orders and 31/50 ECM families (Table 2) in the FungalTraits database that exhaustively summaries all known fungi with primarily ECM lifestyle (Põlme et al. 2020). The most diverse ECM families included *Russulaceae* (140 OTUs), *Thelephoraceae* (94 OTUs), *Sebacinaceae* (63 OTUs), *Inocybaceae* (60 OTUs), *Tuberaceae* (22 OTUs), and *Cortinariaceae* (17 OTUs). Frequent SAP families included *Clavariaceae* (67 OTUs), *Hyaloscyphaceae* (43 OTUs), *Psathyrellaceae* (33 OTUs), *Lyophyllaceae* (17 OTUs), *Omphalotaceae* (17 OTUs), and *Meruliaceae* (17 OTUs). ECM/SAP families of comparatively abundant OTUs were *Pezizaceae* (33 OTUs), *Leotiaceae* (26 OTUs), and *Pyronemataceae* (22 OTUs). However, more than 77 fungal families comprising 21 % of all OTUs were not assigned to the above categories as their members are mostly endophytes, pathogens, or have unknown lifestyles (Fig. 3 and Table S3).

**Fig. 3.**
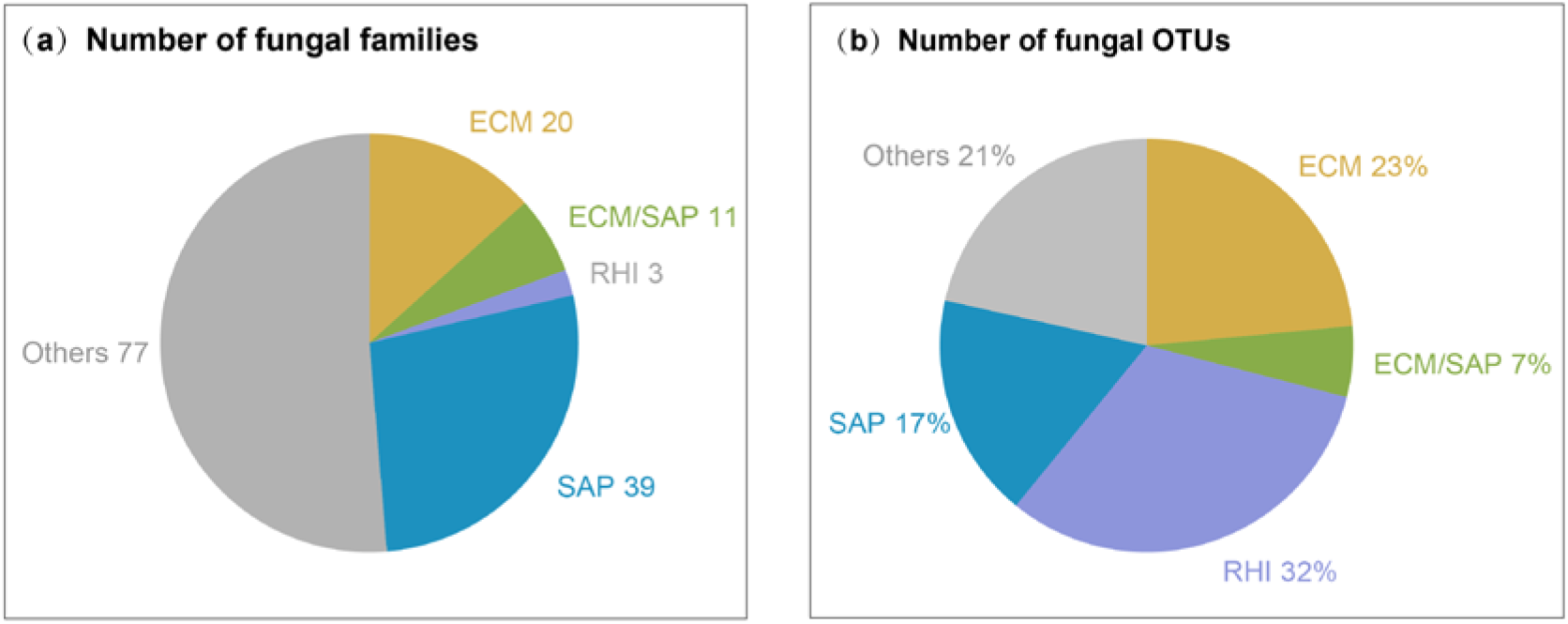
A summary of ecological lifestyles of fungal families and OTUs. (a) The ecological lifestyles of all detected fungal families; (b) the ecological lifestyles of all fungal OTUs. RHI: rhizoctonia fungi; ECM: ectomycorrhizal fungi; SAP: wood/litter-decaying saprotrophic fungi; ECM/SAP: ectomycorrhizal or wood- or litter-decaying saprotrophic fungi; Others: Endophytes, pathogens, unspecified saprotrophs, or unknown fungi.

**Table 2.** The presence of ECM fungal families in the OrM dataset.

| Phylum | Order | Family | Lifestyle coded by the FungalTraits database | Present in OrM dataset |
| --- | --- | --- | --- | --- |
| *Ascomycota* | *Eurotiales* | *Elaphomycetaceae* | ECM | Yes |
| *Ascomycota* | *Gloniales* | *Gloniaceae* | ECM/SAP | – |
| *Ascomycota* | *Helotiales* | *Helotiaceae* | ECM/SAP | Yes |
| *Ascomycota* | *Helotiales* | *Leotiaceae* | ECM/SAP | Yes |
| *Ascomycota* | *Pezizales* | *Discinaceae* | ECM/SAP | Yes |
| *Ascomycota* | *Pezizales* | *Geomoriaceae* | ECM | – |
| *Ascomycota* | *Pezizales* | *Helvellaceae* | ECM | Yes |
| *Ascomycota* | *Pezizales* | *Pezizaceae* | ECM/SAP | Yes |
| *Ascomycota* | *Pezizales* | *Pezizales family*<br>*incertae sedis* | ECM/SAP | – |
| *Ascomycota* | *Pezizales* | *Pulvinulaceae* | ECM/SAP | – |
| *Ascomycota* | *Pezizales* | *Pyronemataceae* | ECM/SAP | Yes |
| *Ascomycota* | *Pezizales* | *Tarzettaceae* | ECM/SAP | – |
| *Ascomycota* | *Pezizales* | *Tuberaceae* | ECM | Yes |
| *Basidiomycota* | *Agaricales* | *Amanitaceae* | ECM/SAP | – |
| *Basidiomycota* | *Agaricales* | *Biannulariaceae* | ECM/SAP | – |
| *Basidiomycota* | *Agaricales* | *Bolbitiaceae* | ECM/SAP | – |
| *Basidiomycota* | *Agaricales* | *Callistosporiaceae* | ECM/SAP | – |
| *Basidiomycota* | *Agaricales* | *Cortinariaceae* | ECM | Yes |
| *Basidiomycota* | *Agaricales* | *Hebelomataceae* | ECM | Yes |
| *Basidiomycota* | *Agaricales* | *Hydnangiaceae* | ECM | Yes |
| *Basidiomycota* | *Agaricales* | *Hygrophoraceae* | ECM/SAP | – |
| *Basidiomycota* | *Agaricales* | *Hymenogastraceae* | ECM/SAP | Yes |
| *Basidiomycota* | *Agaricales* | *Inocybaceae* | ECM | Yes |
| *Basidiomycota* | *Agaricales* | *Lyophyllaceae* | ECM/SAP | Yes |
| *Basidiomycota* | *Agaricales* | *Tricholomataceae* | ECM/SAP | Yes |
| *Basidiomycota* | *Agaricomycetes order*<br>*incertae sedis* | *Agaricomycetes*<br>*family incertae sedis* | ECM/SAP | – |
| *Basidiomycota* | *Atheliales* | *Atheliaceae* | ECM/SAP | Yes |
| *Basidiomycota* | *Boletales* | *Boletaceae* | ECM | – |
| *Basidiomycota* | *Boletales* | *Boletales family*<br>*incertae sedis* | ECM/SAP |  |
| *Basidiomycota* | *Boletales* | *Calostomataceae* | ECM | – |
| *Basidiomycota* | *Boletales* | *Diplocystidiaceae* | ECM | Yes |
| *Basidiomycota* | *Boletales* | *Gomphidiaceae* | ECM | – |
| *Basidiomycota* | *Boletales* | *Gyrodontaceae* | ECM | – |
| *Basidiomycota* | *Boletales* | *Gyroporaceae* | ECM | – |
| *Basidiomycota* | *Boletales* | *Melanogastraceae* | ECM | Yes |
| *Basidiomycota* | *Boletales* | *Paxillaceae* | ECM/SAP | – |
| *Basidiomycota* | *Boletales* | *Pisolithaceae* | ECM | Yes |
| *Basidiomycota* | *Boletales* | *Rhizopogonaceae* | ECM | Yes |
| *Basidiomycota* | *Boletales* | *Sclerodermataceae* | ECM | Yes |
| *Basidiomycota* | *Boletales* | *Serpulaceae* | ECM/SAP | Yes |
| *Basidiomycota* | *Boletales* | *Suillaceae* | ECM | Yes |
| *Basidiomycota* | *Boletales* | *Truncocolumellaceae* | ECM | – |
| *Basidiomycota* | *Cantharellales* | *Cantharellaceae* | ECM | Yes |
| *Basidiomycota* | *Cantharellales* | *Clavulinaceae* | ECM | Yes |
| *Basidiomycota* | *Cantharellales* | *Hydnaceae* | ECM | Yes |
| *Basidiomycota* | *Gomphales* | *Gomphaceae* | ECM/SAP | – |
| *Basidiomycota* | *Hymenochaetales* | *Hymenochaetaceae* | ECM/SAP | Yes |
| *Basidiomycota* | *Hysterangiales* | *Gallaceaceae* | ECM | – |
| *Basidiomycota* | *Hysterangiales* | *Hysterangiaceae* | ECM | – |
| *Basidiomycota* | *Hysterangiales* | *Mesophelliaceae* | ECM | – |
| *Basidiomycota* | *Hysterangiales* | *Phallogastraceae* | ECM | – |
| *Basidiomycota* | *Russulales* | *Albatrellaceae* | ECM | – |
| *Basidiomycota* | *Russulales* | *Russulaceae* | ECM | Yes |
| *Basidiomycota* | *Sebacinales* | *Sebacinaceae* | ECM | Yes |
| *Basidiomycota* | *Thelephorales* | *Bankeraceae* | ECM | Yes |
| *Basidiomycota* | *Thelephorales* | *Thelephoraceae* | ECM | Yes |
| *Basidiomycota* | *Tremellodendropsidales* | *Tremellodendropsida*<br>*ceae* | ECM | – |
| *Mucoromycota* | *Endogonales* | *Densosporaceae* | ECM | – |
| *Mucoromycota* | *Endogonales* | *Endogonaceae* | ECM/SAP | – |

Note: According to the FungalTraits v1.2 database (Põlme et al. 2020), we assign fungal families with the ‘ECM’ lifestyle when the ‘primary lifestyle’ of the family is exclusively recorded as ‘ectomycorrhizal’; we assign fungal families with the ‘ECM/SAP’ lifestyle when the family includes ectomycorrhizal members.

### 3.2 OrM fungi in major orders of *Basidiomycota* and *Ascomycota*

#### 3.2.1 OrM fungi in *Basidiomycota*

##### (1) Cantharellales

The order *Cantharellales* comprised 447 OTUs associating with orchids, yielding the highest number of OTUs of all fungal orders (Fig. 1 and 2, Table S3). Considering the estimated total species number (851 species) of this order by Catalogue of Life (CoL, https://www.catalogueoflife.org/), *Cantharellales* is unquestionably the order with the highest diversity of orchid mycorrhiza forming taxa. Within this order, members were the most abundant in the two ‘rhizoctonia’ families: *Ceratobasidiaceae* and *Tulasnellaceae* (Fig. 4). The prevalence of the two rhizoctonia families has been well reported for orchids (Smith & Read 2008; Dearnaley et al. 2012) and representative strains are found along a wide phylogenetic range of Orchidaceae including the first diverging subfamily Apostasioideae (Yukawa et al. 2009; Wang et al. 2021), in different life stages from protocorm and seedling to adult stages (Rasmussen 2014; Rammitsu et al. 2019), in terrestrial, lithophytic and epiphytic growth forms (Martos et al. 2012; Cevallos et al. 2017; Xing et al. 2019), and across multiple habitats from temperate grasslands to tropical forests at continental and global scales (Shefferson et al. 2007; McCormick & Jacquemyn 2014; Jacquemyn et al. 2017a). Most *Tulasnellaceae* and *Ceratobasidiaceae* fungi are thought to be free-living saprotrophs (Moore & Roberts 2000; Smith & Read 2008; Kohler et al. 2015), but some are also endophytic, pathogenic, and ectomycorrhizal with other land plants (Veldre et al. 2013; Tedersoo & Smith 2013, 2017; Selosse et al. 2018). Orchids obtain nutrients from rhizoctonia fungi either by receiving nutrients through healthy fungal hyphae or by digesting degraded hyphae intracellularly (Dearnaley et al. 2012). Stable isotope cellular imaging has shown that both live and degenerating fungal pelotons formed by *Ceratobasidium* transfer carbon and nitrogen to symbiotic protocorms of *Spiranthes sinensis* (Kuga et al. 2014).

**Fig. 4.**
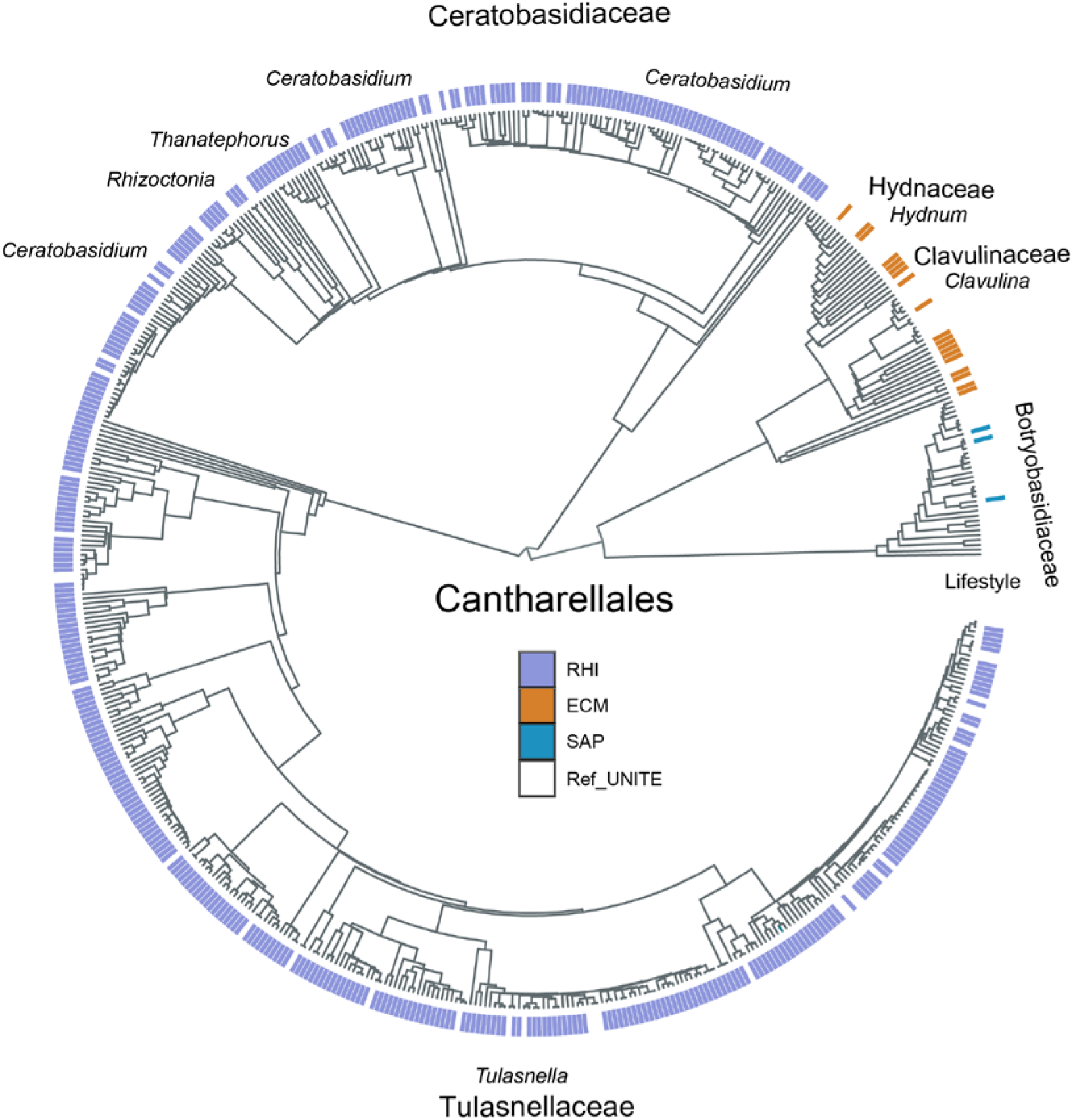
Phylogeny and ecological styles of *Cantharellales*. The ecological lifestyles of fungal OTUs were highlighted in the outer circle: RHI-Rhizotonia fungi in purple, ECM-ectomycorrhizal fungi in orange, SAP-wood/litter-decaying saprotrophic fungi in blue. Reference sequences from the UNITE database were marked as ‘Ref_UNITE’ and are not marked by a color.

Furthermore, three ECM fungal families in *Cantharellales* comprising a small number of OTUs were recorded: *Clavulinaceae* (10 OTUs), *Cantharellaceae* (6 OTUs), and *Hydnaceae* (3 OTUs) (Fig. 4, Table S3). These ectomycorrhizal members have been detected in the roots of several orchid species, such as *Epipactis helleborine* and *Dactylorhiza majalis* (Suetsugu et al. 2017; Schweiger et al. 2018). In addition, saprotrophic *Botryobasidiaceae* (3 OTUs) (Table S3) were reported to associate with *Apostasia wallichii* (Yukawa et al. 2009).

##### (2) Sebacinales

*Sebacinales* is one of the major orders forming orchid mycorrhiza and comprises two subclades with divergent fungal lifestyles: *Sebacinales* group A (*Sebacinaceae*) and group B (*Serendipitaceae*) (Weiß et al. 2004, 2016; Oberwinkler et al. 2013). A total of 243 OTUs of *Sebacinales* were found associating with orchids (Fig. 2, Fig. 5, and Table S3), representing ca. 25 % of the total species of *Sebacinales* recorded in the current UNITE database (Weiß et al. 2016). *Serendipitaceae* contains only one genus *Serendipita* (includes anamorph genus *Piriformospora*) and consists mainly of root endophytes, soil saprotrophs, and ectomycorrhizal taxa (Tedersoo & Smith 2013; Weiß et al. 2016), as well as ericoid mycorrhizal fungi (Selosse et al. 2007; van der Heijden et al. 2015). Members of *Serendipita* form ubiquitously associations with orchids as one of the major groups of rhizoctonia fungi (Dearnaley et al. 2012; Rasmussen et al. 2015), as is also shown by the large number of OTUs recorded in this study (180 OTUs, Fig. 5, and Table S3).

**Fig. 5.**
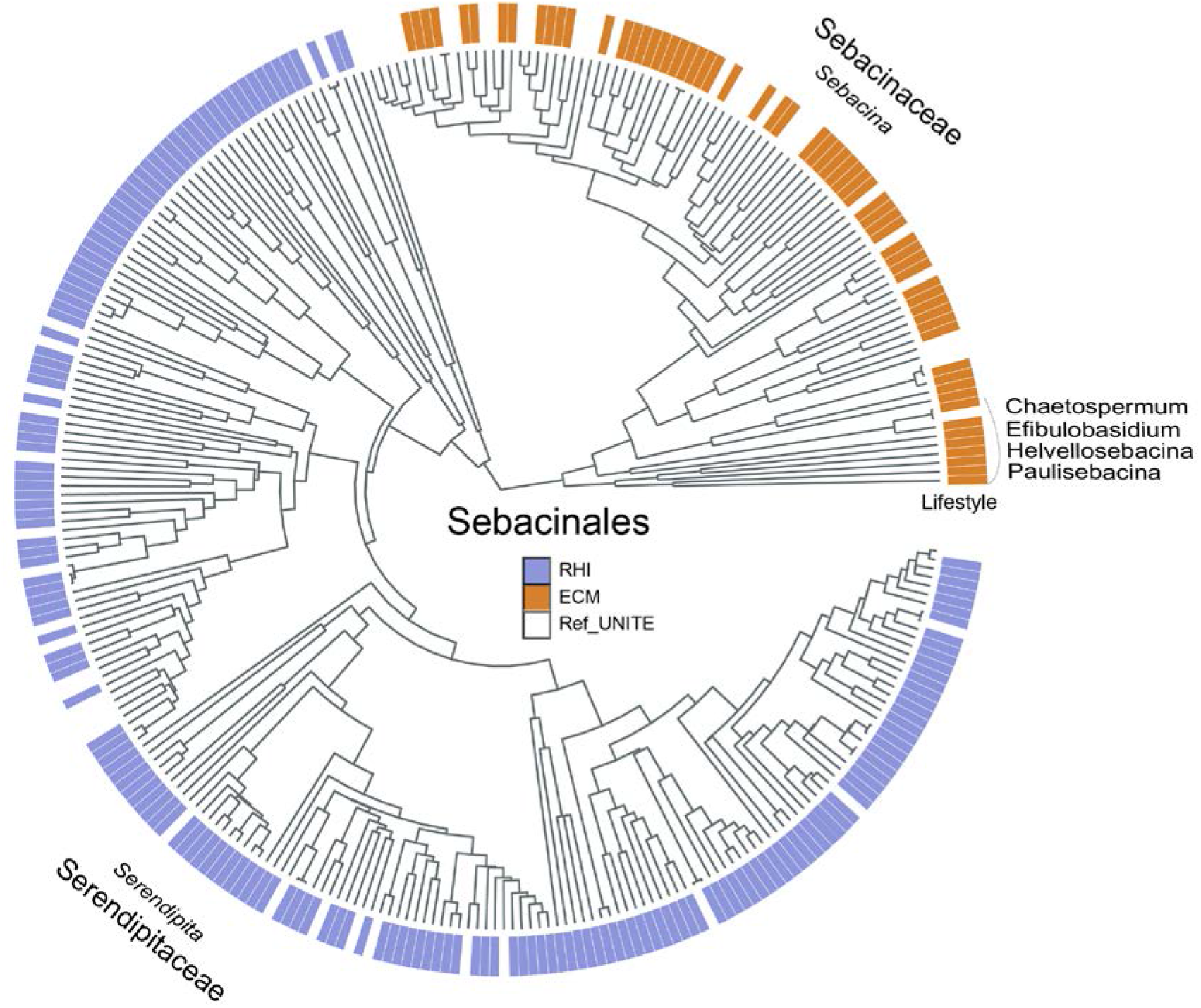
Phylogeny and ecological lifestyles of *Sebacinales*. The ecological lifestyles of fungal OTUs were highlighted in the outer circle: RHI-rhizoctonias in purple, and ECM-ectomycorrhizal fungi in orange. Reference sequences from the UNITE database were marked as ‘Ref_UNITE’ and are not marked by a color.

*Sebacinaceae* mainly contains ectomycorrhizal fungi and this mode of life is regarded as a derived feature, which evolved from endophytic or wood saprotrophic ancestors (Selosse et al. 2009; Weiß et al. 2016). In this study, 63 OTUs belonging to *Sebacinaceae* were found in orchid roots (Fig. 5, Table S3). Interestingly, members of *Sebacinaceae* have mostly been detected in partially and fully mycoheterotrophic orchids that lack the ability to photosynthesize at the adult age, whereas members of *Serendipitaceae* have mainly been found to associate with autotrophic relatives (Taylor et al. 2003; Kennedy et al. 2011; Těšitelová et al. 2015). The symbiotic shift from ‘rhizoctonias’ in *Serendipitaceae* to ectomycorrhizal fungi in *Sebacinaceae* has been suggested to correlate with evolutionary transitions from autotrophy to mycoheterotrophy in orchids (Jacquemyn & Merckx 2019; Wang et al. 2021). This transition is particularly evident in the orchid subtribe Neottieae, which contains the full range of trophic modes from autotrophy via partial to full mycoheterotrophy (Stöckel et al. 2014; Těšitelová et al. 2015; Yagame et al. 2016).

##### (3) Agaricales

*Agaricales*, with at least 24,450 species in the CoL checklist, is the largest order of mushroom-forming fungi (Matheny et al. 2006; Binder et al. 2010; Varga et al. 2019) and is well known to form mycorrhizal interactions with orchids (Rasmussen 2002; Taylor et al. 2002; Smith & Read 2008; Dearnaley et al. 2012). In this study, a total of 296 OTUs were found among 22 families of *Agaricales*, representing the second-highest number of OTUs among all orders (Fig. 2, Fig. 6, and Table S3). *Agaricales* contains multiple ectomycorrhizal members (McLaughlin & Spatafora 2014; Sánchez-García & Matheny 2017; Cao et al. 2021). At least 11 origins of ectomycorrhizal habits have evolved asynchronously in *Agaricales*, resulting in more than 5000 species that can form ectomycorrhiza (Matheny et al. 2006; Rinaldi et al. 2008), not to mention lately recognized ectomycorrhizal lineages in this order (e.g. /porpoloma, /phaeocollybia, /guyanagarika) (Tedersoo et al. 2010; Tedersoo & Smith 2013, 2017). Several ectomycorrhizal families in this order (e.g. *Inocybaceae, Cortinariaceae*, and *Hymenogastraceae*) have been frequently detected in orchid roots using high-throughput sequencing techniques (Jacquemyn et al. 2014), especially for well-studied European terrestrial orchid genera *Epipactis* (Jacquemyn et al. 2016, 2011; Xing et al. 2020) and *Neottia* (Jacquemyn et al. 2015; Těšitelová et al. 2015).

**Fig. 6.**
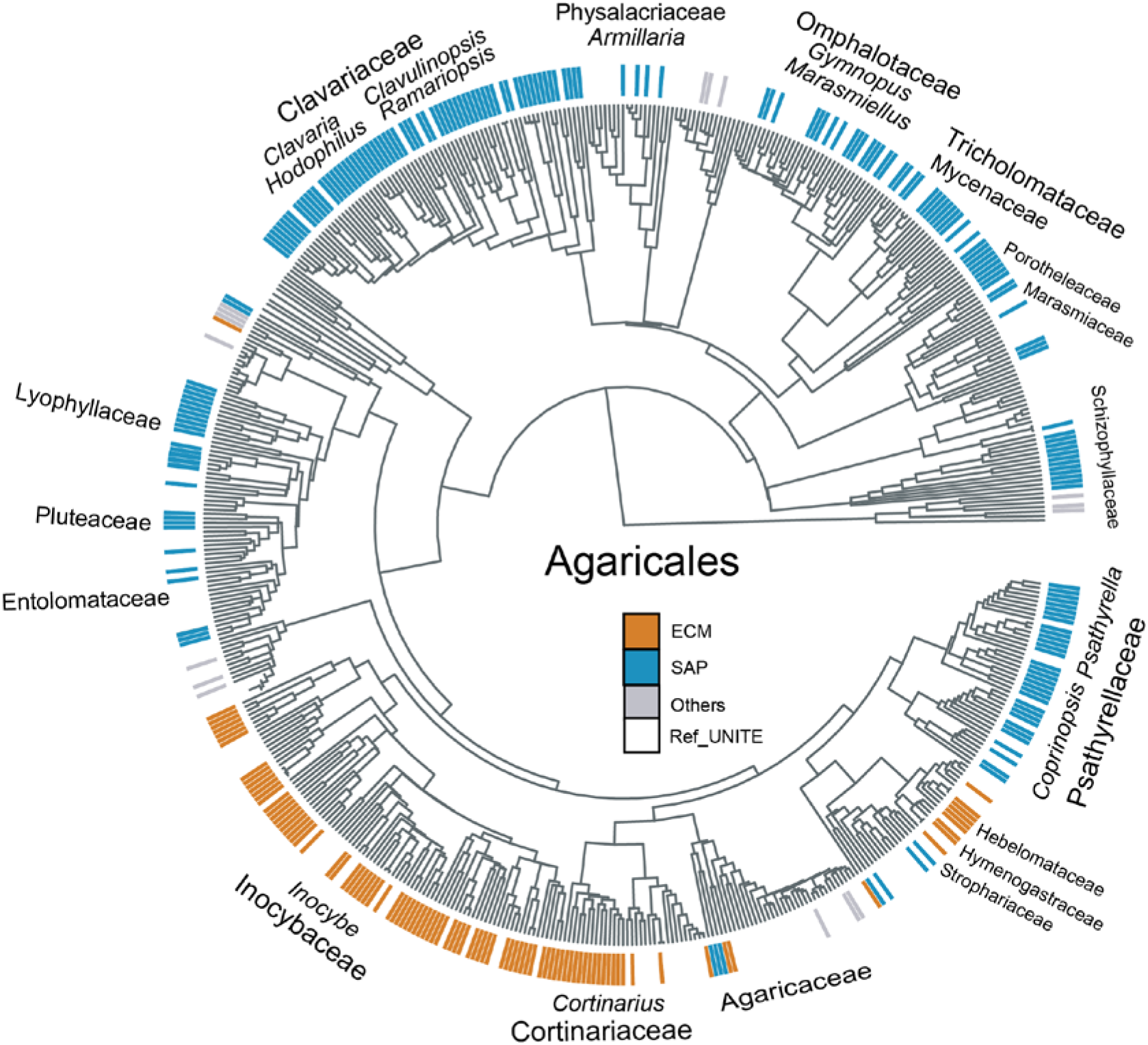
Phylogeny and ecological lifestyles of *Agaricales*. The ecological lifestyles of fungal OTUs were highlighted in the outer circle: ECM-ectomycorrhizal fungi in orange; SAP-wood/litter-decaying saprotrophic fungi in blue; and Others-Endophytic, pathogenic, unspecific saprotrophic, or unknown fungi in gray. Reference sequences from the UNITE database were marked as ‘Ref_UNITE’ and are not marked by a color.

Some saprotrophic members of *Agaricales* have been frequently reported as symbionts with mycoheterotrophic orchids (Ogura-Tsujita et al. 2021). Several saprotrophic families in *Agaricales* (e.g. *Agaricaceae*, *Mycenaceae*, *Marasmiaceae*, *Omphalotaceae*, and *Physalacriaceae*) have been found to associate with a wide range of fully mycoheterotrophic orchids, including orchids from the mycoheterotrophic genera *Gastrodia*, *Galeola*, *Didymoplexis*, *Epipogium*, *Erythrorchis*, *Eulophia*, and *Wullschlaegelia* (Ogura-Tsujita et al. 2021). Litter decaying *Mycena* (*Mycenaceae*) and wood-decaying *Armillaria* (*Physalacriaceae*) are the dominant symbionts of the *Gastrodia* orchid species (Xu & Mu 1990; Xu & Guo 2000; Ogura-Tsujita et al. 2009; Guo et al. 2016). Litter-decaying *Gymnopus* (Omphalotaceae) has also recently been identified as common symbionts with adult individuals of *G. confusoides* within a bamboo forest (Li et al. 2022). Species of the genus *Gastrodia* frequently change fungal partners between the seedling stage and the adult stage (Xu & Mu 1990; Li et al. 2022). Strains of *Mycena* and *Armillaria* have been successfully applied for symbiotic seed germination of fully mycoheterotrophic *Gastrodia* species (Xu & Guo 1989; Park & Lee 2013; Li et al. 2020) and *Cyrtosia septentrionalis* (Umata et al. 2013), as well as several autotrophic species, e.g. *Bletilla striata* (Guo & Xu 1992), *Cymbidium sinense* (Fan et al. 1996), *Anoectochilus roxburghii* (Guo et al. 1997), and *Dendrobium officinale* (Zhang et al. 2012).

##### (4) Russulales

The order *Russulales* comprises at least 3,268 species and ca. 65 % of the total species belongs to the ectomycorrhizal family *Russulaceae* (the CoL checklist). Our results showed that *Russulaceae* contained at least 140 OTUs that associated with orchids and the majority of these was assigned to *Russula* (Fig. 7, Table S3), a speciose genus with more than 1300 species (Bhunjun et al. 2022). Since the first molecular identification of *Russula* as symbionts of *Corallorhiza manculata* (Taylor & Bruns 1997), members of *Russulaceae* have been frequently identified as important mycorrhizal partners of several mycoheterotrophic orchids, including *Corallorhiza* (Taylor & Bruns 1997, 1999; Taylor et al. 2004; Whitridge & Southworth 2005), *Chamaegastrodia* (Pecoraro et al. 2020), *Cymbidium* (Whitridge & Southworth 2005; Motomura et al. 2010; Ogura-Tsujita et al. 2012), *Dipodium* (Dearnaley & Le Brocque 2006), *Erythrorchis* (Dearnaley 2006a), *Epipactis* (Selosse et al. 2004; Těšitelová et al. 2012; Jacquemyn et al. 2016), *Hexalectris* (Kennedy et al. 2011), *Limodorum* (Girlanda et al. 2005), and *Lecanorchis* (Okayama et al. 2012).

**Fig. 7.**
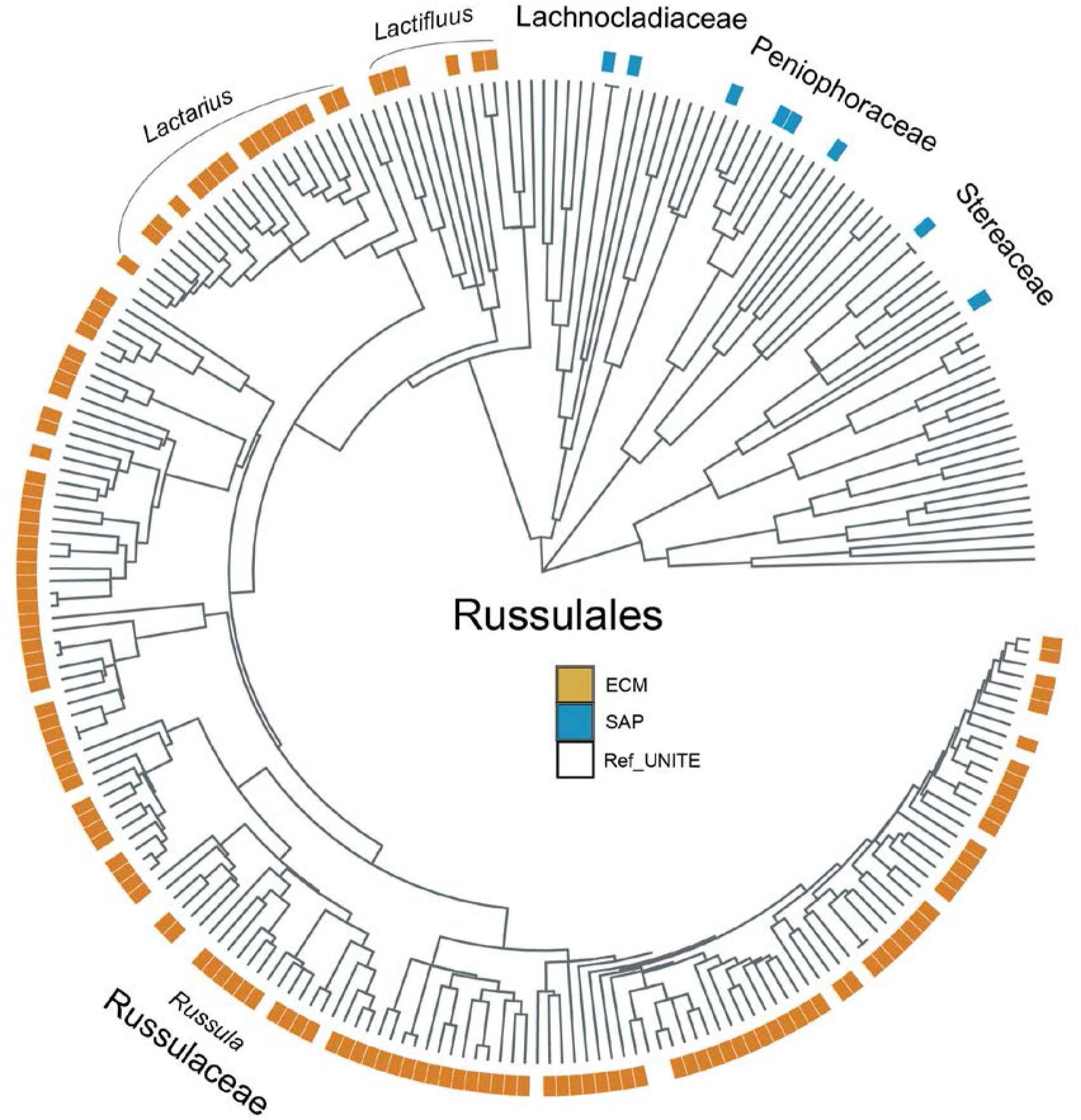
Phylogeny and ecological lifestyles of *Russulales*. The ecological lifestyles of fungal OTUs were highlighted in the outer circle: ECM-ectomycorrhizal fungi in orange; and SAP-wood/litter-decaying saprotrophic fungi in blue. Reference sequences from the UNITE database were marked as ‘Ref_UNITE’ and are not marked by a color.

##### (5) Thelephorales

The order *Thelephorales* contains at least 406 species (the CoL checklist) and most members forming orchid mycorrhizas are ectomycorrhizal (Rasmussen 2002; Taylor et al. 2002; Smith & Read 2008; Dearnaley et al. 2012). In this study, 83 out of 94 OTUs that belong to *Thelephorales* and associate with orchids were assigned to the genus *Tomentella* (*Thelephoraceae*) (Fig. 8, Table S3). *Thelephoraceae* fungi were first identified as symbionts in the mycoheterotrophic orchids *Cephalanthera austinae* and *C. trifida* (Taylor & Bruns 1997; McKendrick et al. 2000b). Since then, shared hyphal connections of *Thelephoraceae* between *C. trifida* and surrounding trees were confirmed by linking the fungal hyphae to roots of *Betula pendula* and *Salix repens* in microcosms and by carbon isotope tracing experiment (McKendrick et al. 2000a). More recently, the participation of *Thelephoraceae* fungi in a triple symbiosis was confirmed between the mycoheterotrophic orchid *C. falcata* and *Quercus serrata* (Fagaceae) in a pot culture experiment (Yagame & Yamato 2013). In addition, the presence of *Thelephoraceae* fungi has been reported not only for mycoheterotrophic orchids across multiple habitats in temperate and tropical regions (Julou et al. 2005; McCormick et al. 2009; Roy et al. 2009; Barrett et al. 2010) but also for green orchids such as *N. ovata* (Těšitelová et al. 2015; Jacquemyn et al. 2015) and *S. spiralis* (Duffy et al. 2019).

**Fig. 8.**
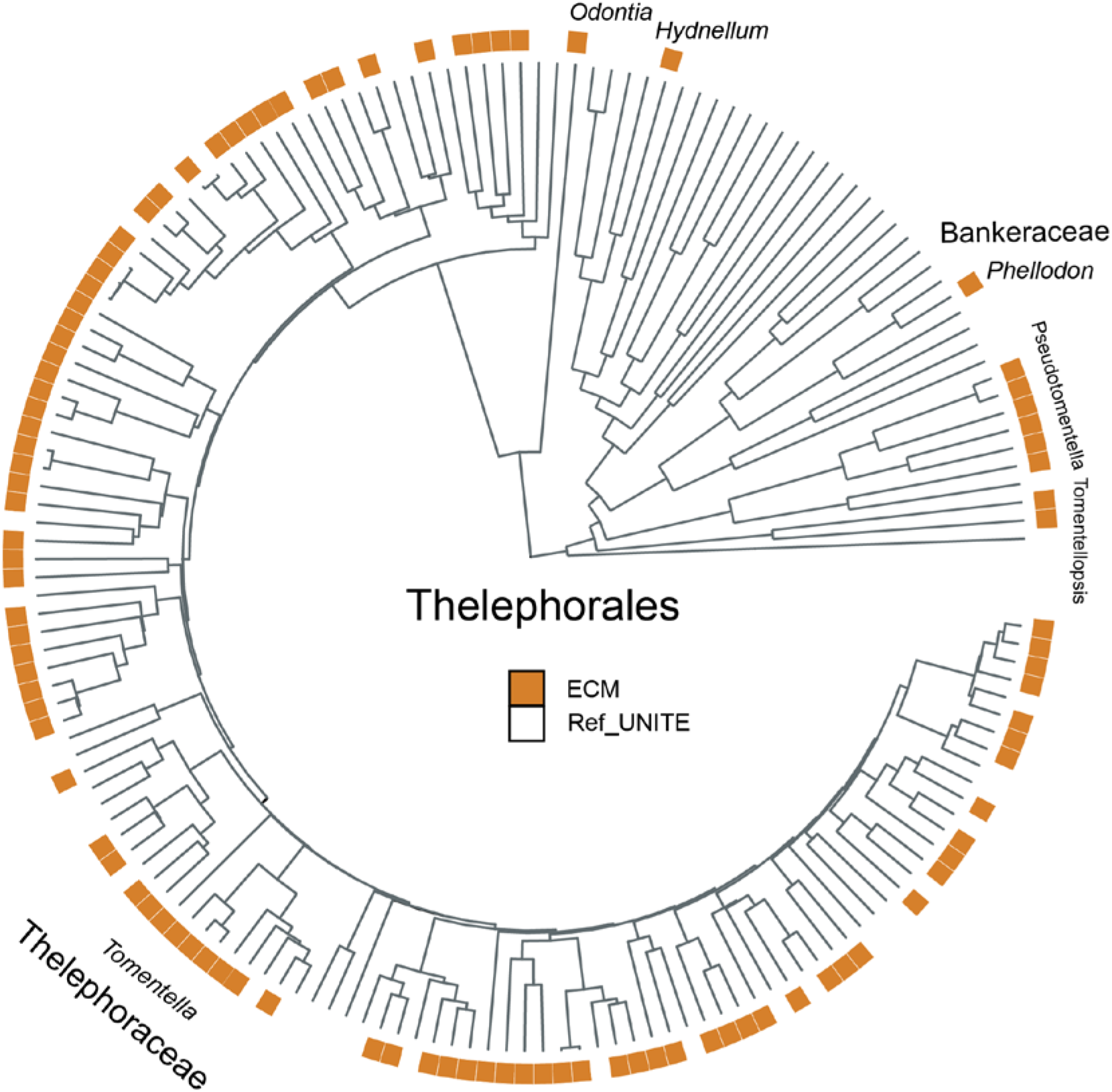
Phylogeny and ecological lifestyles of *Thelephorales*. The ecological lifestyles of fungal OTUs were highlighted in the outer circle: ECM-ectomycorrhizal fungi in orange. Reference sequences from the UNITE database were marked as ‘Ref_UNITE’ and are not marked by a color.

##### (6) Other orders of less prevalent OTUs in Basidiomycota

###### Polyporales

With a total of 3,781 species recorded in the CoL checklist, the order *Polyporales* consists mainly of wood-decaying fungi and some members occasionally have an endophytic or pathogenic lifestyle (Binder et al. 2013; McLaughlin & Spatafora 2014; Martin et al. 2015). We found a total of 32 OTUs of *Polyporales* to associate with orchids and half of them were assigned to the family *Meruliaceae*, which consists mainly of wood saprotrophs (Table S3). Members of *Meruliaceae* have been found as symbionts in the mycoheterotrophic *Erythrorchis* and *Yoania* species based on seed germination experiments (Ogura-Tsujita et al. 2018; Yamashita et al. 2020). In addition, fungi of the family *Polyporaceae* (including *Ganoderma* in previous *Ganodermataceae*) have been detected in the mycoheterotrophic orchids *G. altissima* and *E. ochobiensis* (Umata 1995, 1999).

###### Auriculariales

The order *Auriculariales* comprises several wood-decaying fungi (Weiß & Oberwinkler 2001; McLaughlin & Spatafora 2014). A total of 15 OTUs have been found associating with orchids and these belonged to three distinct families (*Exidiaceae*, *Auriculariaceae*, and *Hyaloriaceae*) (Table S3). For example, *Exidiaceae* species have been detected in the roots of epiphytic and lithophytic *Pleione* species (Qin et al. 2019). *Auricularia polytricha (Auriculariaceae*) has been shown to contribute to the seed germination of the achlorophyllous orchid *E. ochobiensis* (Umata 1997), while unknown *Auriculariales* species have been detected in the protocorms of the autotrophic *Tipularia discolor* (McCormick et al. 2004).

###### Trechisporales

Fourteen OTUs were documented in the family *Hydnodontaceae* in the order *Trechisporales* (Table S3). This family comprises abundant soil-dwelling saprotrophs, some of which (e.g. strains of the type genus *Trechispora*) are possibly root endophytes or even ectomycorrhizal symbionts associated with plant roots (Hibbett et al. 2007; McLaughlin & Spatafora 2014; Vanegas-León et al. 2019). *Trechispora* was found in the protocorms of the mycoheterotrophic *G. nipponica* in an *in-vitro* culture experiment (Shimaoka et al. 2017). Members of *Hydnodontaceae* have also been found to associate with the giant mycoheterotrophic orchid *E. altissima* (Ogura-Tsujita et al. 2018) and several tropical orchids on Réunion Island (Martos et al. 2012).

###### Atractiellales

Members of *Atractiellales*, with 12 OTUs recorded in this study (Table S3), are assumed to be saprotrophic on plant material (Bauer et al. 2006) or endophytic within roots (Oberwinkler et al. 2006; Bonito et al. 2017; Aime et al. 2018). Molecular sequencing and microscopic observations have suggested that *Atractiellales* can engage in a symbiotic association with several tropical and subtropical orchids (Kottke et al. 2010; Cevallos et al. 2018a; Qin et al. 2019; Xing et al. 2019). Moreover, transmission electron microscopy showed that the morphological traits of the hyphal structures formed by *Atractiellales* members closely resemble peloton-like structures formed by typical orchid mycorrhizal fungi (Kottke et al. 2010), indicating their ability to form novel mycorrhizal associations with orchids.

###### Hymenochaetales

Eleven OTUs belonging to the order *Hymenochaetales* were found to associate with orchids and most of them belonged to the family *Hymenochaetaceae* (Table S3). Although most species of this family are wood-decaying saprotrophs, it also comprises ectomycorrhizal genera such as *Coltricia* and *Coltriciella* (Larsson et al. 2006; Tedersoo et al. 2007; McLaughlin & Spatafora 2014). *Coltricia* was reported to colonize the roots of the mycoheterotrophic orchid *E. cassythoides* (Dearnaley 2006b). In addition, the wood-decaying saprotroph *Resinicium* (*Rickenellaceae*) was found in the mycoheterotrophic orchid *G. similis* (Martos et al. 2009).

###### Boletales

While saprotrophy is likely the ancestral state of fungi in the order *Boletales*, the majority of species in this order form ectomycorrhizas with a large variety of conifers and broadleaved trees (Binder & Hibbett 2006; Tedersoo et al. 2010; McLaughlin & Spatafora 2014; Cao et al. 2021). Ten OTUs of this order were found to associate with orchids and mainly belonged to ectomycorrhizal families (Table S3). The ectomycorrhizal species *Scleroderma areolatum* (*Sclerodermataceae*) was found to form peloton-like structures in the roots of *Vanilla* species (González-Chávez et al. 2018), suggesting that it can form mycorrhizal associations with orchids. Besides, saprotrophic fungi of the families *Coniophoraceae* and *Serpulaceae* have been occasionally detected in orchid roots (e.g. *Gymnadenia conopsea*), but their function remains unclear (Těšitelová et al. 2013).

###### Atheliales

Eight OTUs of the order *Atheliales* were found to associate with orchids, all of which belonged to the family *Atheliaceae* (Table S3), the biggest family of the order *Atheliales* (McLaughlin & Spatafora 2014; Cao et al. 2021). This family contains both saprotrophic and ectomycorrhizal fungi (Tedersoo et al. 2010; Wang & Guo 2010; Tedersoo & Smith 2017). Several genera (*Amphinema, Byssocorticium, Piloderma*, and *Tylospora*) have been shown to engage in prevalent symbiotic associations with trees of the Pinaceae and Fagaceae (McLaughlin & Spatafora 2014; Cao et al. 2021). Several members of *Atheliaceae* have been reported as potential mycorrhizal symbionts of orchids (Illyés et al. 2009; Jacquemyn et al. 2010, 2011). In the terrestrial orchid *E. atrorubens*, the ectomycorrhizal fungus *Amphinema* was shown to facilitate seed germination and seedling growth (Bidartondo & Read 2008).

###### Malasseziales

Members of the *Malasseziales* belong to the subphylum *Ustilaginomycotina* and mainly comprise plant pathogens (Begerow et al. 2006). The family *Malasseziaceae* is the only family in the order *Malasseziales* (Cao et al. 2021) and some of its members are well known as inhabitants of human skin (Wu et al. 2015). Members of this family (eight OTUs in total – Table 3) have also been occasionally detected in the roots of orchid species such as *Epipactis* (Těšitelová et al. 2012), *Aphyllorchis* (Roy et al. 2009), and *Spiranthes* (Tondello et al. 2012). However, their mycorrhizal ability and ecological role in orchid roots are unknown and warrant more research.

###### Tremellales

The order *Tremellales* includes a large number of yeasts, saprotrophic and lichen-forming species (Millanes et al. 2011; Liu et al. 2015; Kurtzman & Boekhout 2017). Six OTUs in this order were recovered from the roots of orchid species (Table S3), but their ecological role is unclear (Martos et al. 2012; Hong et al. 2015).

#### 3.2.2 OrM fungi in *Ascomycota*

##### (1) Helotiales

With a total of 6,266 species recorded in the CoL list, many species of the order *Helotiales* are known to be plant pathogens, dark septate endophytes, secondary colonizers of ectomycorrhizal roots, or well-known symbionts for ericoid mycorrhizas (Jumpponen & Trappe 1998; Tedersoo et al. 2009; Kühdorf et al. 2015; Wijayawardene et al. 2017). Moreover, several ectomycorrhizal subclades have recently been reported in this order including /leotia, /helotiales1, /phaeohelotium, /helotiales3, /helotiales4, /helotiales5, /helotiales6 (Tedersoo & Smith 2017). This order contained the highest number of OTUs (104 OTUs) of all *Ascomycota* orders found in the roots of orchids in our dataset (Fig. 2, Table S2). The majority of OTUs were assigned to the family *Hyaloscyphaceae* (43 OTUs) and *Leotiaceae* (26 OTUs), mainly comprising wood/litter saprotrophs (Table S3). Symbiotic culturing and microscopic observations have shown that the hyphae of a dark septate endophytic fungus *Leptodontidium* sp. (*Helotiales incertae sedis*) was able to invade the velamen layer of *D. nobile* roots, penetrate through passage cells and form fungal pelotons in cortex cells (Hou & Guo 2009). Furthermore, *Helotiales* members have been repeatedly detected in the rhizosphere and roots of several orchid genera, including *Anacamptis* (Pecoraro et al. 2018), *Cattleya* (Oliveira et al. 2014), *Cephalanthera* (Julou et al. 2005; Abadie et al. 2006), *Chloraea* (Herrera et al. 2017), *Cymbidium* (Shefferson et al. 2005; Hong et al. 2015), *Cypripedium* (Oja et al. 2015), *Dactylorhiza* (Schiebold et al. 2018), *Epipactis* (Bidartondo et al. 2004; Jacquemyn et al. 2016; Xing et al. 2020), *Goodyera* (Shefferson et al. 2010), *Gymnadenia* (Stark et al. 2009; Xing et al. 2020), *Himantoglossum* (Pecoraro et al. 2013), *Malaxis* (Schiebold et al. 2018), *Liparis* (Schiebold et al. 2018), *Neottia* (Bidartondo et al. 2004; Oja et al. 2015; Těšitelová et al. 2015), *Orchis* (Oja et al. 2015), *Platanthera* (Esposito et al. 2016), *Pleione* (Qin et al. 2019), *Pseudorchis* (Kohout et al. 2013) and *Spiranthes* (Tondello et al. 2012). However, more morphological and physiological evidence is needed to confirm the mycorrhizal status of *Helotiales* fungi in orchids.

##### (2) Pezizales

With a total of 2,788 species in the CoL checklist, the order *Pezizales* mainly contains ectomycorrhizal fungi, wood/litter saprotrophs, plant pathogens, and endophytes (Hansen & Pfister 2006; Wijayawardene et al. 2017; Ekanayaka et al. 2018). The majority of the 92 OTUs belonging to this order that has been observed in the roots of orchids in our dataset (Fig. 2, Table S3) were assigned to three families (*Tuberaceae*, *Pezizaceae*, and *Pyronemataceae*) and comprised almost exclusively ectomycorrhizal fungi. Using molecular identification techniques and electron microscopy, Selosse et al. (2004) showed that fungi from the truffle genus *Tuber* formed peloton-like structures in the roots of *E. microphylla*, representing the first evidence for the symbiotic ability of *Pezizales* with orchids (Selosse et al. 2004). Since then, several other ectomycorrhizal genera within the order *Pezizales*, including *Tuber*, *Peziza*, *Genea, Geopora, Wilcoxina*, and *Trichophaea*, have been recovered in the roots of several orchid species belonging to different genera: *Anacamptis* (Ercole et al. 2015), *Cephalanthera* (Julou et al. 2005), *Corycium* (Waterman et al. 2011), *Cymbidium* (Huang & Zhang 2015), *Epipactis* (Selosse et al. 2004; Bidartondo et al. 2004; May et al. 2020; Xing et al. 2020), *Gymnadenia* (Stark et al. 2009; Těšitelová et al. 2013), *Herminium* (Schiebold et al. 2018), *Limodorum* (Selosse et al. 2004; Girlanda et al. 2005; Těšitelová et al. 2012; Stöckel et al. 2014), and *Pterygodium* (Waterman et al. 2011).

##### (3) Hypocreales

Members of the order *Hypocreales* have several lifestyles, including plant pathogens, soil/litter/wood saprotrophs, and parasites on plants and other organisms (Tedersoo et al. 2009; Wijayawardene et al. 2017; Zhang et al. 2018). Fungi of the family *Nectriaceae* have often been reported as root endophytes of orchids (Bayman & Otero 2007; Tao et al. 2008; Salazar et al. 2020). Here, we recovered 21 OTUs of *Nectriaceae* that have been reported to be associating with orchids (Table S3), representing the largest number of OTUs in the order *Hypocreales*. This family includes numerous important plant pathogens or endophytes (e.g. *Fusarium*) (Summerell et al. 2011). Particularly, several *Fusarium* isolates have been reported to promote seed germination or growth of orchids, including both terrestrial (*Cypripedium* (Vujanovic 2000), *Eulophia* (Johnson et al. 2007), and *Bletilla* (Jiang et al. 2019)) and epiphytic *Dendrobium* (Chen et al. 2010). The earliest reports on the ability of *Fusarium* to stimulate orchid seed germination can be traced back to the 1990s (Vujanovic 2000). The hyphae of *Fusarium oxysporum* have been found to penetrate root hairs and form peloton-like structures within the cortex cells of the roots of *B. striata* (Jiang et al. 2019), which is similar to typical orchid mycorrhizal fungi. In addition, a few isolates of *Trichoderm*a (*Hypocreaceae*) and *Clonostachys* (*Bionectriaceae*) have been used in seed germination experiments. Especially, electron microscopy showed that fungal hyphae of *Clonostachys* were present in living cells of protocorms of the mycoheterotrophic orchid *Pogoniopsis schenckii* (Sisti et al. 2019) despite the absence of peloton formation. Altogether, these studies suggest that fungi of *Hypocreales* may have the potential to be mycorrhizal in orchids and fulfill important ecological functions.

##### (4) Other less prevalent orders in Ascomycota

###### Pleosporales

Species in the order *Pleosporales* can be plant pathogens, endophytes, or wood/litter/soil saprotrophs (Kolařík et al. 2017; Geml 2018). A total of 56 OTUs of this order was found to associate with orchids, with most OTUs belonging to *Phaeosphaeriaceae* and *Pleosporaceae*, two families comprising plant pathogens and wood/litter saprotrophs (Table S3). The potential mycorrhizal status of fungi belonging to this order was demonstrated once based on molecular identification of *Pyrenophora seminiperda (Pleosporaceae*) in isolated pelotons within root cells of *Vanilla* species (González-Chávez et al. 2018).

###### Xylariales

Members of *Xylariales* have diverse ecological lifestyles, including plant pathogens, endophytes, and wood/litter saprotrophs (Wijayawardene et al. 2017; Geml 2018). A total of 43 OTUs associating with orchids were recorded and most of them were assigned to the family *Xylariaceae* (Table S3). Most of these species were reported as endophytes in orchid roots (Dearnaley et al. 2012), especially in roots of epiphytic orchids (Yuan et al. 2009; Cevallos et al. 2018b).

###### Capnodiales

The order *Capnodiales* includes saprotrophic, plant pathogenic, and endophytic species (Wijayawardene et al. 2017; Geml 2018). This study documented 41 OTUs of *Capnodiales* recovered from orchid roots and more than half were assigned to the family *Extremaceae* (Table S3), a family comprising plant pathogens and soil/litter saprotrophs (Wijayawardene et al. 2017). Members in *Capnodiales* have been occasionally detected in orchid roots by molecular sequencing (Těšitelová et al. 2012; Martos et al. 2012; Oliveira et al. 2014; Shah et al. 2018), but their exact ecological role in orchid roots remains unclear.

###### Chaetothyriales

Although most species within the order *Chaetothyriales* are root endophytes, some species act as saprotrophs, endophytes, and ectomycorrhizal fungi (Tedersoo et al. 2009; Wijayawardene et al. 2017; Geml 2018). Fungi of *Chaetothyriales* are also engaged in ericoid mycorrhiza and form intracellular coils within ericoid roots (Watkinson 2016). The majority of OTUs of this order associating with orchids belonged to the family *Herpotrichiellaceae* (27 out of 33 OTUs, Table S3). Members of *Chaetothyriales* were occasionally detected in the rhizosphere of orchid roots (Herrera et al. 2010; Oja et al. 2017) and were found to enhance orchid seedling growth and drought resistance (Liu et al. 2022).

###### Eurotiales

The order *Eurotiales* is mainly comprised of saprotrophs and occasionally ectomycorrhizal fungi (Geml 2018). In this study, a total of 27 OTUs of three *Eurotiales* families were found: apart from one OTU belonging to ectomycorrhizal *Elaphomycetaceae*, the remaining OTUs were assigned to *Aspergillaceae* and *Trichocomaceae*, comprising saprotrophic members (Table S3). Members of *Eurotiales* have only been occasionally isolated from orchid roots or rhizospheres, but so far their mycorrhizal status with orchids has not been verified (Dearnaley & Le Brocque 2006; Oliveira et al. 2014; Hong et al. 2015; Herrera et al. 2017; González-Chávez et al. 2018).

###### Sordariales

The order *Sordariales* mostly consists of plant pathogens, soil saprotrophs, foliar endophytes, and putative ectomycorrhizal fungi (Tedersoo et al. 2009; Wijayawardene et al. 2017; Geml 2018). A total of 23 OTUs were found for three families in this order (*Lasiosphaeriaceae*, *Sordariaceae*, and *Chaetomiaceae*) (Table S3). Members of *Sordariales* have only been occasionally detected in orchid roots (Oliveira et al. 2014; Huang & Zhang 2015), and their mycorrhizal status remains unknown.

###### Glomerellales

The majority of species of *Glomerellales* are plant pathogens, but saprotrophic species also occur in this order (Wijayawardene et al. 2017). The 12 OTUs found associating with orchids were assigned to three families (*Glomerellaceae*, *Plectosphaerellaceae*, and *Australiascaceae*) (Table S3). *Glomerellales* members have rarely been reported as fungal associates of orchids and their mycorrhizal status has not been confirmed (Xing et al. 2011).

###### Saccharomycetales

Twelve OTUs were found in the order *Saccharomycetales* belonging to five distinct families: *Trigonopsidaceae*, *Dipodascaceae*, *Metschnikowiaceae*, *Debaryomycetaceae*, and *Phaffomycetaceae* (Table S3). All of these families contain saprotrophs or endophytes (Wijayawardene et al. 2017). *Saccharomycetales* members have occasionally been detected in orchid species with no observations of their mycorrhizal status with orchids (Těšitelová et al. 2012; Martos et al. 2012).

###### Trichosphaeriales

Eight OTUs belonging to the family *Trichosphaeriaceae* in the order *Trichosphaeriales* were recovered from the roots of orchids (Table S3), but their ecological function remains unclear (Těšitelová et al. 2012; Těšitelová et al. 2015; Ogura-Tsujita et al. 2018; Xing et al. 2019).

###### Diaporthales

Seven OTUs belonging to the order *Diaporthales* were found associating with orchids and these belonged to three distinct families (*Diaporthaceae, Sydowiellaceae*, and *Valsaceae*) (Table S3). Given that most species of this order mainly are plant pathogens or wood/litter saprotrophs (Wijayawardene et al. 2017) and that these fungi have only been rarely detected in orchid roots, their mycorrhizal status is dubious (Bunch et al. 2013; Khamchatra et al. 2016; Herrera et al. 2017; Wang et al. 2017).

###### Chaetosphaeriales

The order *Chaetosphaeriales* mostly consists of wood/litter saprotrophs and endophytes (Geml 2018). We recovered only four OTUs were recovered that were associated with orchids and these belonged to the family *Chaetosphaeriaceae* (Table S3). However, species in *Chaetosphaeriaceae* are only rarely reported as fungal associates of orchids and their mycorrhizal status with orchids has not been examined in detail (Oliveira et al. 2014).

### 3.3 Implications for future studies

#### 3.3.1 A wide range of orchid-associated fungi awaits further examinations of their mycorrhizal status

A large number of fungal OTUs were recorded in this study (Fig. 1 and 2; Table S3) and their taxonomic breadth clearly went beyond the typical rhizoctonia-forming fungi (*Ceratobasidiaceae*, *Tulasnellaceae*, and *Serendipitaceae*) (Smith & Read 2008), confirming earlier summaries of putative orchid mycorrhizal fungi (Dearnaley et al. 2012; van der Heijden et al. 2015; Rasmussen et al. 2015). The identification of a wider taxonomic breadth of orchid-associated fungi is mainly the result of better detection techniques and the increase in sampling intensity of orchid species from diverse phylogenetic clades and ecological niches in the past two decades (Ogura-Tsujita et al. 2021; Li et al. 2021; Wang et al. 2021). It should be noted that this study has only summarized the fungal taxa in *Basidiomycota* and *Ascomycota* that have been frequently detected in ca. 750 out of at least 27,000 orchid species. Thus, it seems certain that more fungal taxa will be detected with the accumulation of studies in the future. Moreover, fungal groups of rare occurrence in orchids may have been overlooked within or outside the two phyla. For example, members of *Mucoromycota* (including previous *Glomeromycota* and *Zygomycota*) have been detected in orchid roots by DNA sequencing despite the uncertainty of their mycorrhizal status with orchids (Cowden & Shefferson 2013; Zhao et al. 2014). In addition, although several general primer sets targeting OrM fungi were used in previous studies, not all fungal groups were amplified by these primer sets, taking the highly variable ITS region of *Tulasnellaceae* as an example (Taylor & McCormick 2008; Waud et al. 2014; Li et al. 2021), indicating a hidden taxonomic diversity of fungal partners of orchids.

According to Rasmussen et al. (2015), the criteria for identifying mycorrhizal fungi of orchid seedlings include peloton isolation, microscopic observation, and symbiotic co-culture experiments. These criteria can also be applied to identify the fungal symbionts of adult orchids. Most fungal DNAs for sequencing were directly extracted from orchid roots with basic root surface sterilizations instead of specific extraction from isolated fungal pelotons (Zhu et al. 2008; Ma et al. 2015). Therefore, it’s largely uncertain whether the fungi extracted from orchid roots are truly mycorrhizal (Rasmussen et al. 2015). Moreover, only a small fraction of studies applied microscopic observations to check the presence and morphology of fungal colonization or conducted symbiotic culture experiments to examine the ecological function of fungal strains (Table S3). Therefore, future studies are highly recommended to combine morphological, physiological, and molecular methods for fungal identification. On the other hand, it’s unlikely that all fungi can be cultured and therefore peloton isolation and subsequent culturing may underestimate the true mycorrhizal diversity in orchid roots.

Despite the uncertainty of their mycorrhizal status, the fungal groups that were frequently detected in the roots of different orchid species and at multiple geographic locations most likely play important ecological roles in the life cycle of orchids. For instance, a dark septate endophyte from Helotiales was found to form peloton-like structures in cortical cells and to promote the growth of *Dendrobium* seedlings (Hou & Guo 2009). The commonly known plant pathogenic fungus *Fusarium* (*Nectriaceae*, *Hypocreales*) has been reported to promote seed germination of several orchid species, including *Cypripedium*, *Platanthera* (Vujanovic 2000), *Bletilla* (Jiang et al. 2019), and *Pogoniopsis* (Sisti et al. 2019). Some endophytic fungi may produce a variety of bioactive compounds and promote orchids’ resistance to biotic and abiotic stresses (Ma et al. 2015). Inoculation of a dark septate endophyte *Exophiala* strain (*Chaetothyriales*) was also found to enhance the drought resistance of *Coelogyne* seedlings (Liu et al. 2022). These studies highlight the potential ecological significance of root endophytes and raise the question of whether they can act as mycorrhizal fungi under certain environmental conditions or in particular life stages of orchids. However, research is still limited on the physiological and ecological functions of putative endophytes for orchids in contrast to much more intensive studies on typical orchid mycorrhizal fungi. The large number of endophytes documented in this study calls for future investigations on their potential mycorrhizal ability, interactions with typical orchid mycorrhizal fungi, and their ecological roles in the ecophysiology of orchids.

Finally, root-associated fungi also receive benefits from the symbiotic relationship with orchids (Dearnaley & Cameron 2017). Green orchids can provide photosynthetic C resources for mycorrhizal fungi to achieve a temporal symbiotic mutualism within orchid lifecycles (Cameron et al. 2006). Although C supply from mycoheterotrophic orchids is limited or not possible, several pioneering studies have already provided evidence that fungi receive other resources (vitamins and ammonia) from orchids (Hijner & Arditti 1973; Cameron et al. 2008), even at early mycoheterotrophic developmental stages of orchids (Fochi et al. 2017). In addition, endophytism (residing in orchid roots) may help to expand the ecological niches of mycorrhizal fungi (Selosse et al. 2018). Since investigations are extremely scarce from the fungal perspective, many of the documented OrM fungi in this study await future in-depth ecophysiological and molecular examinations on their potential benefits from mycorrhizal networks.

#### 3.3.2 The phylogenetic range of OrM fungi serves as an overview for orchid-fungus coevolutionary studies

The evolutionary history of orchid mycorrhiza has fascinated evolutionary biologists for centuries as this type of mycorrhiza exclusively occurs in the highly diversified Orchidaceae family (Rasmussen 1995; Givnish et al. 2015; Rasmussen et al. 2015). Phylogenetic analyses have confirmed the monophyly of Orchidaceae as the sister group of the remainder of Asparagales (Chase et al. 2015; Chomicki et al. 2015; Givnish et al. 2015; Li et al. 2019) and extensive eco-physiological studies have characterized orchid mycorrhizal fungi in all subfamilies and major clades of Orchidaceae (Yukawa et al. 2009; Wang et al. 2021). Thus, it has been suggested that orchid mycorrhizal fungi were already present in the common ancestor of Orchidaceae (Yukawa et al. 2009; Rasmussen 2014) and have possibly evolved from a pre-mycorrhizal status forming intracellular hyphae within orchid root cells. The family-wide ancestral symbiotic state reconstructions provide clues that a diverse fungal community already resided in the roots of the common ancestor of orchids and that multiple symbiotic transitions have taken place in later evolutionary stages (Wang et al. 2021). This is in line with the ‘waiting room’ hypothesis, which states that a variety of fungi reside in orchid roots as non-mycorrhizal endophytes predating the evolutionary transitions to a tight mycorrhizal symbiosis (Selosse et al. 2022).

As shown in this study, we detected a large diversity of fungal lifestyles of orchid-fungal associates including rhizoctonia fungi, ectomycorrhizal, wood-/litter-decaying saprotrophic fungi, and other endophytes (Fig. 1, Table S3). The transitions in fungal lifestyles await future studies that thoroughly trace the evolutionary paths towards particular lifestyles in different fungal clades, as well as with orchid species of distinctive physiological traits, ecological lifeforms, and evolutionary histories. The large-scale phylogenetic framework of orchid-associated fungi that we inferred in this study (Fig. 1; Appendix S4), together with previously published phylogenetic frameworks of the Orchidaceae family (Chase et al. 2015; Chomicki et al. 2015; Givnish et al. 2015; Li et al. 2019; Wang et al. 2021), can provide clues for selecting interesting clades of orchids and their mycorrhizal fungi for future ecological and co-evolutionary studies of orchid-fungal interactions.

#### 3.3.3 Conservation aspects

Orchid mycorrhizal fungi are obligate for seed germination and seedling development for nearly all orchids (Rasmussen 1995; Smith & Read 2008; Dearnaley et al. 2012). Identifying the core symbiont fungi is critical for the conservation and restoration of rare and threatened orchid populations (Dearnaley 2007). For instance, both *ex-situ* and *in-situ* symbiotic germination methods have been successfully applied in orchid conservation for decades mainly by inoculations of rhizoctonia fungal strains (Rasmussen 1995; Brundrett et al. 2003; Kottke & Nebel 2005; Batty et al. 2006). Apart from typical rhizoctonia fungi, we documented a large number of ectomycorrhizal and wood- or litter-decaying saprotrophic fungi (Table S3). The saprotrophic decomposers living on wood debris or leaf litter have been shown to provide sufficient nutrients to fully mycoheterotrophic orchids in forests (Leake 1994; Ogura-Tsujita et al. 2009, 2021; Martos et al. 2009; Selosse et al. 2010). Owing to their free-living properties, several saprotrophic fungi have been successfully cultured and *in/ex-situ* cultures have been established to facilitate the conservation of endangered orchid species (Xu and Guo 2000; Yagame et al. 2007; Shimaoka et al. 2017).

Because the ectomycorrhizal fungi establish a tripartite mycorrhizal network including surrounding vegetation and orchids, the symbiotic culture of orchids associated with ectomycorrhizal fungi is challenging, as both an autotrophic plant and a particular locally distributed ectomycorrhizal fungus are required for the successful germination and development of these orchid species (McKendrick et al. 2000a, 2002; Bougoure et al. 2010). Furthermore, orchid-associated fungal communities can vary among regional habitats (Oja et al. 2015; Jacquemyn et al. 2016) and even between micro-climates of tree barks and forest floors in the tropics (Martos et al. 2012). Therefore, revealing ecological lifestyles and environmental niches of symbiotic fungi will contribute to understanding the distribution and population dynamics of orchids and developing new strategies for their conservation. We hope that this overview of the diversity of orchid root-associated fungi will stimulate future in-depth ecological studies on orchid-fungal interactions and conservation practices.

## 4 Conclusion

Based on a newly compiled molecular dataset of orchid-associated fungi, we revisited and summarized a broad taxonomic range of orchid-fungal associates belonging to at least 150 families of 28 orders in *Basidiomycota* and *Ascomycota*. Apart from their taxonomic diversity, these fungal taxa were found to exhibit diverse ecological lifestyles, ranging from the typical rhizoctonia-forming fungi to ectomycorrhizal fungi, wood- or litter-decaying saprotrophic fungi, and other endophytic, saprotrophic, and pathogenic fungi. However, the mycorrhizal ability and ecological lifestyle of most fungal taxa still await verification. Nevertheless, we hope that the newly reconstructed phylogenetic framework of orchid-associated fungi can contribute to future studies on orchid mycorrhiza from an orchid-fungus co-evolutionary perspective. We conclude that OrM fungi are probably the most speciose group of mycorrhizal fungi that deserves more intensive studies on their taxonomic identity, ecological lifestyle, and evolutionary history.

## Acknowledgments

We sincerely thank for the funding provided by the China Scholarship Council (Grant No. 201804910634) and the Ecology Fund of the Royal Netherlands Academy of Arts and Sciences (KNAWWF/807/19039).

## Author contributions

V.S.F.T.M. initiated and supervised the project. D.W. created the dataset. D.W. and J.L. performed the analyses with support and guidance from J.N., S.I.F.G., H.J., and V.S.F.T.M. All authors contributed to the writing of the manuscript.

## Conflict of Interest

The authors declare no conflict of interest.

## Data Availability

The datasets analyzed during the current study are available in in open figshare database (doi: 10.6084/m9.figshare.21158875).

## Supplementary Information

**Appendix S1** Dataset of orchid-associated fungal sequences after filtering.fasta

**Appendix S2** Dataset of orchid-associated fungal sequences assigned with order names.fasta

**Appendix S3** Dataset of orchid-associated fungal OTUs.fasta

**Appendix S4** Alignments and trees.zip

**Appendix S5** Supertree phylogeny of orchid root-associated fungi in Newick format.txt

**Table S1** Fungal OTUs, and DOIs (SHs) of the corresponding UNITE species hypothesis.xlsx

**Table S2** Number of sequences, OTUs, divergence time, and outgroups of 28 fungal orders.xlsx

**Table S3** Number of OTUs and ecological lifestyles, and orchid mycorrhizal status of 150 fungal family.xlsx

## Notes

https://doi.org/10.6084/m9.figshare.21158875

### Competing Interest Statement

The authors have declared no competing interest.

## References

Abadie JC, Püttsepp Ü, Gebauer G, et al (2006) *Cephalanthera longifolia* (Neottieae, Orchidaceae) is mixotrophic: a comparative study between green and nonphotosynthetic individuals. Can J Bot 84:1462–1477. https://doi.org/10.1139/b06-101

Aime, Urbina, Liber, et al (2018) Two new endophytic Atractiellomycetes, *Atractidochium hillariae* and *Proceropycnis hameedii*. Mycologia 110:136–146. https://doi.org/10.1080/00275514.2018.1446650

Barrett CF, Freudenstein J V., Lee Taylor D, Kõljalg U (2010) Rangewide analysis of fungal associations in the fully mycoheterotrophic *Corallorhiza striata* complex (Orchidaceae) reveals extreme specificity on ectomycorrhizal *Tomentella* (Thelephoraceae) across North America. Am J Bot 97:628–643. https://doi.org/10.3732/ajb.0900230

Batty AL, Brundrett MC, Dixon KW, Sivasithamparam K (2006) New methods to improve symbiotic propagation of temperate terrestrial orchid seedlings from axenic culture to soil. Aust J Bot 54:367. https://doi.org/10.1071/BT04023

Bauer R, Begerow D, Sampaio JP, et al (2006) The simple-septate basidiomycetes: A synopsis. Mycol Prog. https://doi.org/10.1007/s11557-006-0502-0

Bayman P, Otero JT (2007) Microbial endophytes of orchid roots. In: Microbial Root Endophytes. Springer Berlin Heidelberg, Berlin, Heidelberg, pp 153–177

Begerow D, Stoll M, Bauer R (2006) A phylogenetic hypothesis of Ustilaginomycotina based on multiple gene analyses and morphological data. Mycologia 98:906–916. https://doi.org/10.3852/mycologia.98.6.906

Bhunjun CS, Niskanen T, Suwannarach N, et al (2022) The numbers of fungi: are the most speciose genera truly diverse? Fungal Divers in press: https://doi.org/10.1007/s13225-022-00501-4

Bidartondo MI, Burghardt B, Gebauer G, et al (2004) Changing partners in the dark: isotopic and molecular evidence of ectomycorrhizal liaisons between forest orchids and trees. Proceedings Biol Sci 271:1799–806. https://doi.org/10.1098/rspb.2004.2807

Bidartondo MI, Read DJ (2008) Fungal specificity bottlenecks during orchid germination and development. Mol Ecol 17:3707–3716

Bidartondo MI, Read DJ, Trappe JM, et al (2011) The dawn of symbiosis between plants and fungi. Biol Lett 7:574–577. https://doi.org/10.1098/rsbl.2010.1203

Binder M, Hibbett DS (2006) Molecular systematics and biological diversification of Boletales. Mycologia 98:971–981. https://doi.org/10.3852/mycologia.98.6.971

Binder M, Justo A, Riley R, et al (2013) Phylogenetic and phylogenomic overview of the Polyporales. Mycologia 105:1350–1373. https://doi.org/10.3852/13-003

Binder M, Larsson KH, Matheny PB, Hibbett DS (2010) Amylocorticiales ord. nov. and Jaapiales ord. nov.: early diverging clades of Agaricomycetidae dominated by corticioid forms. Mycologia 102:865–880. https://doi.org/10.3852/09-288

Blaxter M, Mann J, Chapman T, et al (2005). Defining operational taxonomic units using DNA barcode data. Philosophical Transactions of the Royal Society B: Biological Sciences, 360: 1935–1943. https://doi.org/10.1098/rstb.2005.1725

Bonito G, Hameed K, Toome-Heller M, et al (2017) *Atractiella rhizophila*, sp. nov., an endorrhizal fungus isolated from the Populus root microbiome. Mycologia 109:18–26. https://doi.org/10.1080/00275514.2016.1271689

Bougoure JJ, Brundrett MC, Grierson PF (2010) Carbon and nitrogen supply to the underground orchid, *Rhizanthella gardneri*. New Phytol 186:947–956. https://doi.org/10.1111/j.1469-8137.2010.03246.x

Brundrett MC (2009) Mycorrhizal associations and other means of nutrition of vascular plants: understanding the global diversity of host plants by resolving conflicting information and developing reliable means of diagnosis. Plant Soil 320:37–77. https://doi.org/10.1007/s11104-008-9877-9

Brundrett MC, Scade A, Batty AL, et al (2003) Development of *in situ* and *ex situ* seed baiting techniques to detect mycorrhizal fungi from terrestrial orchid habitats. Mycol Res 107:1210–1220. https://doi.org/10.1017/S0953756203008463

Brundrett MC, Tedersoo L (2018) Evolutionary history of mycorrhizal symbioses and global host plant diversity. New Phytol 220:1108–1115. https://doi.org/10.1111/nph.14976

Bruns TD, Szaro TM, Gardes M, et al (1998) A sequence database for the identification of ectomycorrhizal basidiomycetes by phylogenetic analysis. Mol Ecol 7:257–272. https://doi.org/10.1046/j.1365-294X.1998.00337.x

Bunch WD, Cowden CC, Wurzburger N, Shefferson RP (2013) Geography and soil chemistry drive the distribution of fungal associations in lady’s slipper orchid, *Cypripedium acaule*. Botany 91:850–856. https://doi.org/10.1139/cjb-2013-0079

Cameron DD, Johnson I, Read DJ, Leake JR (2008) Giving and receiving: measuring the carbon cost of mycorrhizas in the green orchid, *Goodyera repens*. New Phytol 180:176–184. https://doi.org/10.1111/j.1469-8137.2008.02533.x

Cameron DD, Leake JR, Read DJ (2006) Mutualistic mycorrhiza in orchids: evidence from plant–fungus carbon and nitrogen transfers in the green-leaved terrestrial orchid *Goodyera repens*. New Phytol 171:405–416. https://doi.org/10.1111/j.1469-8137.2006.01767.x

Cao B, Haelewaters D, Schoutteten N, et al (2021) Delimiting species in Basidiomycota: a review. Fungal Divers 109:181–237. https://doi.org/10.1007/s13225-021-00479-5

Cevallos S, Declerck S, Suárez JP (2018a) *In situ* orchid seedling-trap experiment shows few keystone and many randomly associated mycorrhizal fungal species during early plant colonization. Front Plant Sci 9:. https://doi.org/10.3389/fpls.2018.01664

Cevallos S, Herrera P, Sánchez-Rodríguez A, et al (2018b) Untangling factors that drive community composition of root associated fungal endophytes of Neotropical epiphytic orchids. Fungal Ecol. https://doi.org/10.1016/j.funeco.2018.05.002

Cevallos S, Sánchez-Rodríguez A, Decock C, et al (2017) Are there keystone mycorrhizal fungi associated to tropical epiphytic orchids? Mycorrhiza 27:225–232. https://doi.org/10.1007/s00572-016-0746-8

Chase MW, Cameron KM, Freudenstein J V., et al (2015) An updated classification of Orchidaceae. Bot J Linn Soc 177:151–174. https://doi.org/10.1111/boj.12234

Chen XM, Dong HL, Hu KX, et al (2010) Diversity and antimicrobial and plant-growth-promoting activities of endophytic fungi in *Dendrobium loddigesii* Rolfe. J Plant Growth Regul 29:328–337. https://doi.org/10.1007/s00344-010-9139-y

Chomicki G, Bidel LPR, Ming F, et al (2015) The velamen protects photosynthetic orchid roots against UV-B damage, and a large dated phylogeny implies multiple gains and losses of this function during the Cenozoic. New Phytol 205:1330–1341. https://doi.org/10.1111/nph.13106

Clements MA, Muir H, Cribb PJ (1986) A preliminary report on the symbiotic germination of european terrestrial orchids. Kew Bull 41:437. https://doi.org/10.2307/4102957

Comandini O, Rinaldi AC, Kuyper TW (2012) M Easuring and E Stimating E Ctomycorrhizal. In: Marcela Pagano (ed) Mycorrhiza: Occurrence in Natural and Restored Environments. Nova Science Publishers, pp 165–200

Cowden CC, Shefferson RP (2013) Diversity of root-associated fungi of mature *Habenaria radiata* and *Epipactis thunbergii* colonizing manmade wetlands in Hiroshima Prefecture, Japan. Mycoscience 54:327–334. https://doi.org/10.1016/j.myc.2012.12.001

Dearnaley J (2006a) The fungal endophytes of *Erythrorchis cassythoides* – is this orchid saprophytic or parasitic? Australas Mycol 25:51–57

Dearnaley JDW (2006b) Molecular identification of fungal endophytes in australian myco-heterotrophic orchids. In: 8th International Mycological Congress (IMC8)

Dearnaley JDW (2007) Further advances in orchid mycorrhizal research. Mycorrhiza 17:475–486. https://doi.org/10.1007/s00572-007-0138-1

Dearnaley JDW, Cameron DD (2017) Nitrogen transport in the orchid mycorrhizal symbiosis – further evidence for a mutualistic association. New Phytol 213:10–12. https://doi.org/10.1111/nph.14357

Dearnaley JDW, Le Brocque AF (2006) Molecular identification of the primary root fungal endophytes of *Dipodium hamiltonianum* (Orchidaceae). Aust J Bot 54:487. https://doi.org/10.1071/BT05149

Dearnaley JDW, Martos F, Selosse M-A (2012) Orchid mycorrhizas: molecular ecology, physiology, evolution and conservation aspects. In: Fungal Associations. Springer Berlin Heidelberg, Berlin, Heidelberg, pp 207–230

Dressler RL, Rasmussen HN (1996) Terrestrial Orchids: From Seed to Mycotrophic Plant. Syst Bot 21:625. https://doi.org/10.2307/2419622

Duffy KJ, Waud M, Schatz B, et al (2019) Latitudinal variation in mycorrhizal diversity associated with a European orchid. J Biogeogr 46:968–980. https://doi.org/10.1111/jbi.13548

Edgar RC (2010) Search and clustering orders of magnitude faster than BLAST. Bioinformatics 26:2460–2461. https://doi.org/10.1093/bioinformatics/btq461

Edgar RC (2004) MUSCLE: multiple sequence alignment with high accuracy and high throughput. Nucleic Acids Res 32:1792–1797. https://doi.org/10.1093/nar/gkh340

Ekanayaka AH, Hyde KD, Jones EBG, Zhao Q (2018) Taxonomy and phylogeny of operculate discomycetes: Pezizomycetes. Fungal Divers 90:161–243. https://doi.org/10.1007/s13225-018-0402-z

Ercole E, Adamo M, Rodda M, et al (2015) Temporal variation in mycorrhizal diversity and carbon and nitrogen stable isotope abundance in the wintergreen meadow orchid *Anacamptis morio*. New Phytol 205:1308–1319. https://doi.org/10.1111/nph.13109

Esposito F, Jacquemyn H, Waud M, Tyteca D (2016) Mycorrhizal fungal diversity and community composition in two closely related *Platanthera* (Orchidaceae) species. PLoS One 11:e0164108. https://doi.org/10.1371/journal.pone.0164108

Fan L, Guo S, Cao W, et al (1996) Isolation, culture, identification and biological activity of *Mycena orchidicola* sp. nov. in Cymbidium sinense (Orchidaceae). Acta Mycol Sin 15:251–255

Feijen FAA, Vos RA, Nuytinck J, Merckx VSFT (2018) Evolutionary dynamics of mycorrhizal symbiosis in land plant diversification. Sci Rep 8:10698. https://doi.org/10.1038/s41598-018-28920-x

Fochi V, Chitarra W, Kohler A, et al (2017) Fungal and plant gene expression in the Tulasnella calospora–Serapias vomeracea symbiosis provides clues about nitrogen pathways in orchid mycorrhizas. New Phytol 213:365–379. https://doi.org/10.1111/nph.14279

Frank AB, Trappe JM (2005) On the nutritional dependence of certain trees on root symbiosis with belowground fungi (an English translation of A.B. Frank’s classic paper of 1885). Mycorrhiza 15:267–75. https://doi.org/10.1007/s00572-004-0329-y

Gardes M, and Bruns TD (1993) ITS primers with enhanced specificity for basidiomycetes – application to the identification of mycorrhizae and rusts. Mol. Ecol. 2:113–118. https://doi.org/10.1111/j.1365-294x.1993.tb00005.x

Geml J (2018) Landscape-level DNA metabarcoding study in the Pannonian forests reveals differential effects of slope aspect on taxonomic and functional groups of fungi. bioRxiv

Girlanda M, Selosse MA, Cafasso D, et al (2005) Inefficient photosynthesis in the Mediterranean orchid *Limodorum abortivum* is mirrored by specific association to ectomycorrhizal Russulaceae. Mol Ecol 15:491–504. https://doi.org/10.1111/j.1365-294X.2005.02770.x

Givnish TJ, Spalink D, Ames M, et al (2015) Orchid phylogenomics and multiple drivers of their extraordinary diversification. Proc R Soc B Biol Sci 282:20151553. https://doi.org/10.1098/rspb.2015.1553

González-Chávez M del CA, Torres-Cruz TJ, Sánchez SA, et al (2018) Microscopic characterization of orchid mycorrhizal fungi: *Scleroderma* as a putative novel orchid mycorrhizal fungus of *Vanilla* in different crop systems. Mycorrhiza 28:147–157. https://doi.org/10.1007/s00572-017-0808-6

Guo S-X, Fan L, Cao W-Q, et al (1997) *Mycena anoectochila* sp. nov. isolated from mycorrhizal roots of Anoectochilus roxburghii from Xishuangbanna, China. Mycologia 89:952. https://doi.org/10.2307/3761116

Guo S, Xu J (1992) The relation between the seed germination and seedling development of *Bletilla striata* and *Mycena osmundicola* etc. fungi. Zhongguo Yi Xue Ke Xue Yuan Xue Bao 14:51–4

Guo T, Wang HC, Xue WQ, et al (2016) Phylogenetic analyses of *Armillaria* reveal at least 15 phylogenetic lineages in China, seven of which are associated with cultivated *Gastrodia elata*. PLoS One. https://doi.org/10.1371/journal.pone.0154794

Hansen K, Pfister DH (2006) Systematics of the Pezizomycetes – the operculate discomycetes. Mycologia 98:1029–1040. https://doi.org/10.3852/mycologia.98.6.1029

He M-Q, Zhao R-L, Hyde KD, et al (2019) Notes, outline and divergence times of Basidiomycota. Fungal Divers 99:105–367. https://doi.org/10.1007/s13225-019-00435-4

Herrera H, Valadares R, Contreras D, et al (2017) Mycorrhizal compatibility and symbiotic seed germination of orchids from the Coastal Range and Andes in south central Chile. Mycorrhiza 27:175–188. https://doi.org/10.1007/s00572-016-0733-0

Herrera P, Suárez JP, Kottke I (2010) Orchids keep the ascomycetes outside: a highly diverse group of ascomycetes colonizing the velamen of epiphytic orchids from a tropical mountain rainforest in Southern Ecuador. Mycology 1:262–268. https://doi.org/10.1080/21501203.2010.526645

Hibbett DS, Binder M, Bischoff JF, et al (2007) A higher-level phylogenetic classification of the Fungi. Mycol Res 111:509–547. https://doi.org/10.1016/j.mycres.2007.03.004

Hijner JA, Arditti J (1973) Orchid Mycorrhiza: Vitamin production and requirements by the symbionts. Am J Bot 60:829. https://doi.org/10.2307/2441176

Hong JW, Suh H, Kim OH, Lee NS (2015) Molecular identification of mycorrhizae of *Cymbidium kanran* (Orchidaceae) on Jeju Island, Korea. Mycobiology 43:475–480. https://doi.org/10.5941/MYCO.2015.43.4.475

Hongsanan S, Maharachchikumbura SSN, Hyde KD, et al (2017) An updated phylogeny of Sordariomycetes based on phylogenetic and molecular clock evidence. Fungal Divers 84:25–41. https://doi.org/10.1007/s13225-017-0384-2

Horton TR, Bruns TD (2001) The molecular revolution in ectomycorrhizal ecology: peeking into the black-box. Mol Ecol 10:1855–1871. https://doi.org/10.1046/j.0962-1083.2001.01333.x

Hou X-Q, Guo S-X (2009) Interaction between a dark septate endophytic isolate from *Dendrobium* sp. and roots of D. nobile seedlings. J Integr Plant Biol 51:374–381. https://doi.org/10.1111/j.1744-7909.2008.00777.x

Hoysted GA, Kowal J, Jacob A, et al (2018) A mycorrhizal revolution. Curr Opin Plant Biol 44:1–6. https://doi.org/10.1016/j.pbi.2017.12.004

Huang F, Zhang C (2015) Diversity, host-and habitat-preferences on the fungi communities from the roots of *Cymbidium* spp. at two sites in China. J Anim Plant Sci 25:270–277

Illyés Z, Halász K, Rudnóy S, et al (2009) Changes in the diversity of the mycorrhizal fungi of orchids as a function of the water supply of the habitat. J Appl Bot Food Qual 83:28–36

Jacquemyn H, Brys R, Merckx VSFT, et al (2014) Coexisting orchid species have distinct mycorrhizal communities and display strong spatial segregation. New Phytol 202:616–627. https://doi.org/10.1111/nph.12640

Jacquemyn H, Brys R, Waud M, et al (2021) Mycorrhizal communities and isotope signatures in two partially mycoheterotrophic orchids. Front Plant Sci 12:1–9. https://doi.org/10.3389/fpls.2021.618140

Jacquemyn H, Duffy KJ, Selosse M-A (2017) Biogeography of orchid mycorrhizas. In: Biogeography of Mycorrhizal Symbiosis. pp 159–177

Jacquemyn H, Honnay O, Cammue BPA, et al (2010) Low specificity and nested subset structure characterize mycorrhizal associations in five closely related species of the genus *Orchis*. Mol Ecol 19:4086–4095. https://doi.org/10.1111/j.1365-294X.2010.04785.x

Jacquemyn H, Merckx V, Brys R, et al (2011) Analysis of network architecture reveals phylogenetic constraints on mycorrhizal specificity in the genus *Orchis* (Orchidaceae). New Phytol 192:518–528. https://doi.org/10.1111/j.1469-8137.2011.03796.x

Jacquemyn H, Merckx VSFT (2019) Mycorrhizal symbioses and the evolution of trophic modes in plants. J Ecol 107:1567–1581. https://doi.org/10.1111/1365-2745.13165

Jacquemyn H, Waud M, Lievens B, Brys R (2016) Differences in mycorrhizal communities between *Epipactis palustris*, *E. helleborine* and its presumed sister species *E. neerlandica*. Ann Bot 118:105–114. https://doi.org/10.1093/aob/mcw015

Jacquemyn H, Waud M, Merckx VSFT, et al (2015) Mycorrhizal diversity, seed germination and long-term changes in population size across nine populations of the terrestrial orchid *Neottia ovata*. Mol Ecol 24:3269–3280. https://doi.org/10.1111/mec.13236

Ji B, Bever JD (2012) Mycorrhizal Ecology. In: Ecology. Oxford University Press

Jiang J, Zhang K, Cheng S, et al (2019) *Fusarium oxysporum* KB-3 from *Bletilla striata*: an orchid mycorrhizal fungus. Mycorrhiza 29:531–540. https://doi.org/10.1007/s00572-019-00904-3

Johnson TR, Stewart SL, Dutra D, et al (2007) Asymbiotic and symbiotic seed germination of *Eulophia alta* (Orchidaceae) – preliminary evidence for the symbiotic culture advantage. Plant Cell Tissue Organ Cult 90:313–323. https://doi.org/10.1007/s11240-007-9270-z

Julou T, Burghardt B, Gebauer G, et al (2005) Mixotrophy in orchids: insights from a comparative study of green individuals and nonphotosynthetic individuals of *Cephalanthera damasonium*. New Phytol 166:639–653. https://doi.org/10.1111/j.1469-8137.2005.01364.x

Jumpponen A, Trappe JM (1998) Dark septate endophytes: A review of facultative biotrophic root-colonizing fungi. New Phytol 140:295–310. https://doi.org/10.1046/j.1469-8137.1998.00265.x

Kennedy AH, Taylor DL, Watson LE (2011) Mycorrhizal specificity in the fully mycoheterotrophic *Hexalectris* Raf. (Orchidaceae: Epidendroideae). Mol Ecol. https://doi.org/10.1111/j.1365-294X.2011.05000.x

Khamchatra N, Dixon KW, Tantiwiwat S, Piapukiew J (2016) Symbiotic seed germination of an endangered epiphytic slipper orchid, *Paphiopedilum villosum* (Lindl.) Stein. from Thailand. South African J Bot. https://doi.org/10.1016/j.sajb.2015.11.012

Kohler A, Kuo A, Nagy LG, et al (2015) Convergent losses of decay mechanisms and rapid turnover of symbiosis genes in mycorrhizal mutualists. Nat Genet 47:410–415. https://doi.org/10.1038/ng.3223

Kohout P (2017) Biogeography of ericoid mycorrhiza. In: Biogeography of Mycorrhizal Symbiosis. pp 179–193

Kohout P, Těšitelová T, Roy M, et al (2013) A diverse fungal community associated with *Pseudorchis albida* (Orchidaceae) roots. Fungal Ecol 6:50–64. https://doi.org/10.1016/j.funeco.2012.08.005

Kolařík M, Spakowicz DJ, Gazis R, et al (2017) *Biatriospora* (Ascomycota: Pleosporales) is an ecologically diverse genus including facultative marine fungi and endophytes with biotechnological potential. Plant Syst Evol 303:35–50. https://doi.org/10.1007/s00606-016-1350-2

Kottke I, Nebel M (2005) The evolution of mycorrhiza-like associations in liverworts: An update. New Phytol 167:330–334. https://doi.org/10.1111/j.1469-8137.2005.01471.x

Kottke I, Suárez JP, Herrera P, et al (2010) Atractiellomycetes belonging to the ‘rust’ lineage (Pucciniomycotina) form mycorrhizae with terrestrial and epiphytic neotropical orchids. Proc R Soc B Biol Sci 277:1289–1298. https://doi.org/10.1098/rspb.2009.1884

Kuga Y, Sakamoto N, Yurimoto H (2014) Stable isotope cellular imaging reveals that both live and degenerating fungal pelotons transfer carbon and nitrogen to orchid protocorms. New Phytol 202:594–605. https://doi.org/10.1111/nph.12700

Kühdorf K, Münzenberger B, Begerow D, et al (2015) *Leotia cf. lubrica* forms arbutoid mycorrhiza with *Comarostaphylis arbutoides* (Ericaceae). Mycorrhiza 25:109–120. https://doi.org/10.1007/s00572-014-0590-7

Kurtzman CP, Boekhout T (2017) Yeasts as distinct life forms of fungi. In: Yeasts in Natural Ecosystems: Ecology. Springer International Publishing, Cham, pp 1–37

Larsson KH, Parmasto E, Fischer M, et al (2006) Hymenochaetales: a molecular phylogeny for the hymenochaetoid clade. Mycologia 98:926–936. https://doi.org/10.3852/mycologia.98.6.926

Leake JR (1994) The biology of myco-heterotrophic (saprophytic) plants. New Phytol 127:171–216. https://doi.org/DOI10.1111/j.1469-8137.1994.tb04272.x

Lee YI, Yang CK, Gebauer G (2015) The importance of associations with saprotrophic non-Rhizoctonia fungi among fully mycoheterotrophic orchids is currently under-estimated: Novel evidence from sub-tropical Asia. Ann Bot 116:423–435. https://doi.org/10.1093/aob/mcv085

Li T, Yang W, Wu S, et al (2021) Progress and prospects of mycorrhizal fungal diversity in orchids. Front Plant Sci 12:. https://doi.org/10.3389/fpls.2021.646325

Li YX, Li ZH, Schuiteman A, et al (2019) Phylogenomics of Orchidaceae based on plastid and mitochondrial genomes. Mol Phylogenet Evol 139:106540. https://doi.org/10.1016/j.ympev.2019.106540

Li YY, Boeraeve M, Cho YH, et al (2022) Mycorrhizal switching and the role of fungal abundance in seed germination in a fully mycoheterotrophic orchid, *Gastrodia confusoides*. Front Plant Sci 12:1–12. https://doi.org/10.3389/fpls.2021.775290

Li YY, Guo SX, Lee YI (2020) Ultrastructural changes during the symbiotic seed germination of *Gastrodia elata* with fungi, with emphasis on the fungal colonization region. Bot Stud 61:4. https://doi.org/10.1186/s40529-019-0280-z

Liu XZ, Wang QM, Göker M, et al (2015) Towards an integrated phylogenetic classification of the Tremellomycetes. Stud Mycol 81:85–147. https://doi.org/10.1016/j.simyco.2015.12.001

Liu N, Jacquemyn H, Liu Q, et al (2022). Effects of a dark septate fungal endophyte on the growth and physiological response of seedlings to drought in an epiphytic orchid. Frontiers in Microbiology, 2590. https://doi.org/10.3389/fmicb.202

Ma X, Kang J, Nontachaiyapoom S, et al (2015) Non-mycorrhizal endophytic fungi from orchids. Curr Sci 109:72–87. https://doi.org/10.18520/cs/v109/i1/72-87

Mao H, Wang H (2019) Resolution of deep divergence of club fungi (phylum Basidiomycota). Synth Syst Biotechnol 4:225–231. https://doi.org/10.1016/j.synbio.2019.12.001

Martin R, Gazis R, Skaltsas D, et al (2015) Unexpected diversity of basidiomycetous endophytes in sapwood and leaves of *Hevea*. Mycologia 107:284–297. https://doi.org/10.3852/14-206

Martos F, Dulormne M, Pailler T, et al (2009) Independent recruitment of saprotrophic fungi as mycorrhizal partners by tropical achlorophyllous orchids. New Phytol 184:668–681. https://doi.org/10.1111/j.1469-8137.2009.02987.x

Martos F, Munoz F, Pailler T, et al (2012) The role of epiphytism in architecture and evolutionary constraint within mycorrhizal networks of tropical orchids. Mol Ecol 21:5098–109. https://doi.org/10.1111/j.1365-294X.2012.05692.x

Matheny PB, Curtis JM, Hofstetter V, et al (2006) Major clades of Agaricales: a multilocus phylogenetic overview. Mycologia 98:982–995. https://doi.org/10.3852/mycologia.98.6.982

May M, Jąkalski M, Novotná A, et al (2020) Three-year pot culture of *Epipactis helleborine* reveals autotrophic survival, without mycorrhizal networks, in a mixotrophic species. Mycorrhiza 30:51–61. https://doi.org/10.1007/s00572-020-00932-4

McCormick MK, Whigham DF, O’Neill J (2004) Mycorrhizal diversity in photosynthetic terrestrial orchids. New Phytol 163:425–438. https://doi.org/10.1111/j.1469-8137.2004.01114.x

McCormick MK, Whigham DF, O’Neill JP, et al (2009) Abundance and distribution of *Corallorhiza odontorhiza* reflect variations in climate and ectomycorrhizae. Ecol Monogr 79:619–635. https://doi.org/10.1890/08-0729.1

McKendrick SL, Leake JR, Read DJ (2000a) Symbiotic germination and development of myco-heterotrophic plants in nature: Transfer of carbon from ectomycorrhizal *Salix repens* and *Betula pendula* to the orchid *Corallorhiza trifida* through shared hyphal connections. New Phytol. https://doi.org/10.1046/j.1469-8137.2000.00592.x

McKendrick SL, Leake JR, taylor DL, Read DJ (2000b) Symbiotic germination and development of myco-heterotrophic plants in nature: ontogeny of *Corallorhiza trifida* and characterization of its mycorrhizal fungi. New Phytol 145:523–537. https://doi.org/10.1046/j.1469-8137.2000.00603.x

McKendrick SL, Leake JR, Taylor DL, Read DJ (2002) Symbiotic germination and development of the myco-heterotrophic orchid *Neottia nidus-avis* in nature and its requirement for locally distributed *Sebacina* spp. New Phytol 154:233–247. https://doi.org/10.1046/j.1469-8137.2002.00372.x

McLaughlin DJ, Spatafora JW (2014) Systematics and evolution: Part A: Second edition. Springer Berlin Heidelberg. https://doi.org/10.1007/978-3-642-55318-9

Merckx VSFT (2013) Mycoheterotrophy: The Biology of Plants Living on Fungi. Springer New York Springer New York Heidelberg Dordrecht London. http://doi.org/10.1007/978-1-4614-5209-6

Millanes AM, Diederich P, Ekman S, Wedin M (2011) Phylogeny and character evolution in the jelly fungi (Tremellomycetes, Basidiomycota, Fungi). Mol Phylogenet Evol. https://doi.org/10.1016/j.ympev.2011.05.014

Miller MA, Pfeiffer W, Schwartz T (2010) Creating the CIPRES Science Gateway for inference of large phylogenetic trees. In: 2010 Gateway Computing Environments Workshop, GCE 2010

Moore RT, Roberts P (2000) Rhizoctonia-forming fungi, a taxonomic guide. Kew Bull 55:252. https://doi.org/10.2307/4117793

Motomura H, Selosse M-A, Martos F, et al (2010) Mycoheterotrophy evolved from mixotrophic ancestors: evidence in *Cymbidium* (Orchidaceae). Ann Bot 106:573–581. https://doi.org/10.1093/aob/mcq156

Mujica I, Fernanda P, Jakalski M, et al (2020) Soil P reduces mycorrhizal colonization while favors fungal pathogens: observational and experimental evidence in Bipinnula (Orchidaceae). 1–12. https://doi.org/10.1093/femsec/fiaa178

Nguyen NH, Song Z, Bates ST, et al (2016) FUNGuild: an open annotation tool for parsing fungal community datasets by ecological guild. Fungal Ecol 20:241–248. https://doi.org/10.1016/j.funeco.2015.06.006

Nilsson RH, Larsson K-H, Taylor AFS, et al (2019) The UNITE database for molecular identification of fungi: handling dark taxa and parallel taxonomic classifications. Nucleic Acids Res 47: 259–264. https://doi.org/10.1093/nar/gky1022

Oberwinkler F, Kirschner R, Arenal F, et al (2006) Two new pycnidial members of the Atractiellales: *Basidiopycnis hyalina* and *Proceropycnis pinicola*. Mycologia 98:637–649. https://doi.org/10.3852/mycologia.98.4.637

Oberwinkler F, Riess K, Bauer R, et al (2013) Enigmatic Sebacinales. Mycol Prog 12:1–27. https://doi.org/10.1007/s11557-012-0880-4

Ogura-Tsujita Y, Gebauer G, Hashimoto T, et al (2009) Evidence for novel and specialized mycorrhizal parasitism: the orchid *Gastrodia confusa* gains carbon from saprotrophic *Mycena*. Proc R Soc B Biol Sci 276:761–767. https://doi.org/10.1098/rspb.2008.1225

Ogura-Tsujita Y, Gebauer G, Xu H, et al (2018) The giant mycoheterotrophic orchid *Erythrorchis altissima* is associated mainly with a divergent set of wood-decaying fungi. Mol Ecol 27:1324–1337. https://doi.org/10.1111/mec.14524

Ogura-Tsujita Y, Yokoyama J, Miyoshi K, Yukawa T (2012) Shifts in mycorrhizal fungi during the evolution of autotrophy to mycoheterotrophy in *Cymbidium* (Orchidaceae). Am J Bot 99:1158–1176. https://doi.org/10.3732/ajb.1100464

Ogura-Tsujita Y, Yukawa T, Kinoshita A (2021) Evolutionary histories and mycorrhizal associations of mycoheterotrophic plants dependent on saprotrophic fungi. J Plant Res 134:19–41. https://doi.org/10.1007/s10265-020-01244-6

Oja J, Kohout P, Tedersoo L, et al (2015) Temporal patterns of orchid mycorrhizal fungi in meadows and forests as revealed by 454 pyrosequencing. New Phytol 205:1608–1618. https://doi.org/10.1111/nph.13223

Oja J, Vahtra J, Bahram M, et al (2017) Local-scale spatial structure and community composition of orchid mycorrhizal fungi in semi-natural grasslands. Mycorrhiza 27:355–367. https://doi.org/10.1007/s00572-016-0755-7

Okayama M, Yamato M, Yagame T, Iwase K (2012) Mycorrhizal diversity and specificity in *Lecanorchis* (Orchidaceae). Mycorrhiza 22:545–553. https://doi.org/10.1007/s00572-012-0429-z

Oliveira SF, Bocayuva MF, Veloso TGR, et al (2014) Endophytic and mycorrhizal fungi associated with roots of endangered native orchids from the Atlantic Forest, Brazil. Mycorrhiza 24:55–64. https://doi.org/10.1007/s00572-013-0512-0

Öpik M, Vanatoa A, Vanatoa E, et al (2010) The online database MaarjAM reveals global and ecosystemic distribution patterns in arbuscular mycorrhizal fungi (Glomeromycota). New Phytol 188:223–241. https://doi.org/10.1111/j.1469-8137.2010.03334.x

Paradis E, Schliep K (2019) ape 5.0: an environment for modern phylogenetics and evolutionary analyses in R. Bioinformatics 35:526–528. https://doi.org/10.1093/bioinformatics/bty633

Park EJ, Lee WY (2013) In vitro symbiotic germination of myco-heterotrophic *Gastrodia elata* by *Mycena* species. Plant Biotechnol Rep 7:185–191. https://doi.org/10.1007/s11816-012-0248-x

Pecoraro L, Caruso T, Cai L, et al (2018) Fungal networks and orchid distribution: new insights from above- and below-ground analyses of fungal communities. IMA Fungus 9:1–11. https://doi.org/10.5598/imafungus.2018.09.01.01

Pecoraro L, Girlanda M, Kull T, et al (2013) Fungi from the roots of the terrestrial photosynthetic orchid *Himantoglossum adriaticum*. Plant Ecol Evol 146:145–152. https://doi.org/10.5091/plecevo.2013.782

Pecoraro L, Wang X, Venturella G, et al (2020) Molecular evidence supports simultaneous association of the achlorophyllous orchid *Chamaegastrodia inverta* with ectomycorrhizal Ceratobasidiaceae and Russulaceae. BMC Microbiol 20:236. https://doi.org/10.1186/s12866-020-01906-4

Põlme S, Abarenkov K, Henrik Nilsson R, et al (2020) FungalTraits: a user-friendly traits database of fungi and fungus-like stramenopiles. Fungal Divers 105:1–16. https://doi.org/10.1007/s13225-020-00466-2

Prieto M, Wedin M (2013) Dating the Diversification of the Major Lineages of Ascomycota (Fungi). PLoS One 8:e65576. https://doi.org/10.1371/journal.pone.0065576

Qin J, Zhang W, Ge Z-W, Zhang S-B (2019) Molecular identifications uncover diverse fungal symbionts of *Pleione* (Orchidaceae). Fungal Ecol 37:19–29. https://doi.org/10.1016/j.funeco.2018.10.003

Rammitsu K, Yagame T, Yamashita Y, et al (2019) A leafless epiphytic orchid, *Taeniophyllum glandulosum* Blume (Orchidaceae), is specifically associated with the Ceratobasidiaceae family of basidiomycetous fungi. Mycorrhiza 29:159–166. https://doi.org/10.1007/s00572-019-00881-7

Rasmussen HN (2014) Seedling mycorrhiza: A discussion of origin and evolution in Orchidaceae. Bot J Linn Soc 175:313–327. https://doi.org/10.1111/boj.12170

Rasmussen HN (2002) Recent developments in the study of orchid mycorrhiza. In: Plant and Soil. pp 149–163

Rasmussen HN (1995) Terrestrial orchids: from seed to mycotrophic plant. Cambridge University Press

Rasmussen HN, Dixon KW, Jersáková J, Těšitelová T (2015) Germination and seedling establishment in orchids: a complex of requirements. Ann Bot 116:391–402. https://doi.org/10.1093/aob/mcv087

Read DJ (1991) Mycorrhizas in ecosystems. Experientia 47:376–391. https://doi.org/10.1007/BF01972080

Rinaldi AC, Comandini O, Kuyper TW (2008) Ectomycorrhizal fungal diversity: Separating the wheat from the chaff. Fungal Divers 33:1–45

Roy M, Watthana S, Stier A, et al (2009) Two mycoheterotrophic orchids from Thailand tropical dipterocarpacean forests associate with a broad diversity of ectomycorrhizal fungi. BMC Biol 7:51. https://doi.org/10.1186/1741-7007-7-51

Salazar JM, Pomavilla M, Pollard AT, et al (2020) Endophytic fungi associated with roots of epiphytic orchids in two Andean forests in Southern Ecuador and their role in germination. Lankesteriana 37–47. https://doi.org/10.15517/lank.v20i1.41157

Sánchez-García M, Matheny PB (2017) Is the switch to an ectomycorrhizal state an evolutionary key innovation in mushroom-forming fungi? A case study in the Tricholomatineae (Agaricales). Evolution 71:51–65. https://doi.org/10.1111/evo.13099

Sayers EW, O’Sullivan C, Karsch-Mizrachi I (2022) Using GenBank and SRA. In Plant Bioinformatics (pp. 1–25). Humana, New York, NY.

Schiebold JMI, Bidartondo MI, Karasch P, et al (2017) You are what you get from your fungi: nitrogen stable isotope patterns in *Epipactis* species. Ann Bot 119:1085–1095. https://doi.org/10.1093/aob/mcw265

Schiebold JMI, Bidartondo MI, Lenhard F, et al (2018) Exploiting mycorrhizas in broad daylight: Partial mycoheterotrophy is a common nutritional strategy in meadow orchids. J Ecol 106:168–178. https://doi.org/10.1111/1365-2745.12831

Schweiger JMI, Bidartondo MI, Gebauer G (2018) Stable isotope signatures of underground seedlings reveal the organic matter gained by adult orchids from mycorrhizal fungi. Funct Ecol 32:870–881. https://doi.org/10.1111/1365-2435.13042

Selosse M-A, Dubois M-P, Alvarez N (2009) Do Sebacinales commonly associate with plant roots as endophytes? Mycol Res 113:1062–1069. https://doi.org/10.1016/j.mycres.2009.07.004

Selosse M-A, Faccio A, Scappaticci G, Bonfante P (2004) Chlorophyllous and achlorophyllous specimens of *Epipactis microphylla* (Neottieae, Orchidaceae) are associated with ectomycorrhizal septomycetes, including truffles. Microb Ecol 47:. https://doi.org/10.1007/s00248-003-2034-3

Selosse M-A, Schneider-Maunoury L, Martos F (2018) Time to re-think fungal ecology? Fungal ecological niches are often prejudged. New Phytol 217:968–972. https://doi.org/10.1111/nph.14983

Selosse M-A, Setaro S, Glatard F, et al (2007) Sebacinales are common mycorrhizal associates of Ericaceae. New Phytol 174:864–878. https://doi.org/10.1111/j.1469-8137.2007.02064.x

Selosse M, Petrolli R, Mujica MI, et al (2022) The Waiting Room Hypothesis revisited by orchids: were orchid mycorrhizal fungi recruited among root endophytes? Ann Bot 129:259–270. https://doi.org/10.1093/aob/mcab134

Selosse MA, Martos F, Perry B, et al (2010) Saprotrophic fungal symbionts in tropical achlorophyllous orchids. Plant Signal Behav 5:349–353. https://doi.org/10.4161/psb.5.4.10791

Shah S, Shrestha R, Maharjan S, et al (2018) Isolation and characterization of plant growth-promoting endophytic fungi from the roots of *Dendrobium moniliforme*. Plants 8:5. https://doi.org/10.3390/plants8010005

Shefferson RP, Cowden CC, McCormick MK, et al (2010) Evolution of host breadth in broad interactions: mycorrhizal specificity in East Asian and North American rattlesnake plantains *(Goodyera* spp.) and their fungal hosts. Mol Ecol 19:3008–17. https://doi.org/10.1111/j.1365-294X.2010.04693.x

Shefferson RP, Taylor DL, Weiß M, et al (2007) The evolutionary history of mycorrhizal specificity among lady’s slipper orchids. Evolution 61:1380–1390. https://doi.org/10.1111/j.1558-5646.2007.00112.x

Shefferson RP, Weiss M, Kull T, Taylor DL (2005) High specificity generally characterizes mycorrhizal association in rare lady’s slipper orchids, genus *Cypripedium*. Mol Ecol 14:613–26. https://doi.org/10.1111/j.1365-294X.2005.02424.x

Shimaoka C, Fukunaga H, Inagaki S, Sawa S (2017) Artificial cultivation system for *Gastrodia* spp. and Identification of associated mycorrhizal fungi. Int J Biol 9:27. https://doi.org/10.5539/ijb.v9n4p27

Sisti LS, Flores-Borges DNA, Andrade SAL de, et al (2019) The role of non-mycorrhizal fungi in germination of the mycoheterotrophic orchid *Pogoniopsis schenckii* Cogn. Front Plant Sci 10:1589. https://doi.org/10.3389/fpls.2019.01589

Smith S, Read D (2008) Mycorrhizal Symbiosis. Elsevier

Smith SA, O’Meara BC (2012) TreePL: Divergence time estimation using penalized likelihood for large phylogenies. Bioinformatics. https://doi.org/10.1093/bioinformatics/bts492

Stamatakis A (2014) RAxML version 8: a tool for phylogenetic analysis and post-analysis of large phylogenies. Bioinformatics 30:1312–1313. https://doi.org/10.1093/bioinformatics/btu033

Stark C, Babik W, Durka W (2009) Fungi from the roots of the common terrestrial orchid *Gymnadenia conopsea*. Mycol Res 113:952–959. https://doi.org/10.1016/j.mycres.2009.05.002

Stöckel M, Těšitelová T, Jersáková J, et al (2014) Carbon and nitrogen gain during the growth of orchid seedlings in nature. New Phytol 202:606–615. https://doi.org/10.1111/nph.12688

Suetsugu K, Yamato M, Miura C, et al (2017) Comparison of green and albino individuals of the partially mycoheterotrophic orchid *Epipactis helleborine* on molecular identities of mycorrhizal fungi, nutritional modes and gene expression in mycorrhizal roots. Mol Ecol. https://doi.org/10.1111/mec.14021

Summerell BA, Leslie JF, Liew ECY, et al (2011) *Fusarium* species associated with plants in Australia. Fungal Divers 46:1–27. https://doi.org/10.1007/s13225-010-0075-8

Tao G, Liu ZY, Hyde KD, et al (2008) Whole rDNA analysis reveals novel and endophytic fungi in *Bletilla ochracea* (Orchidaceae). Fungal Divers 33:101–112

Taylor DL, Bruns TD (1997) Independent, specialized invasions of ectomycorrhizal mutualism by two nonphotosynthetic orchids. Proc Natl Acad Sci U S A 94:4510–5. https://doi.org/10.1073/pnas.94.9.4510

Taylor DL, Bruns TD (1999) Population, habitat and genetic correlates of mycorrhizal specialization in the “cheating” orchids *Corallorhiza maculata* and *C. mertensiana*. Mol Ecol 8:1719–1732. https://doi.org/10.1046/j.1365-294x.1999.00760.x

Taylor DL, Bruns TD, Hodges SA (2004) Evidence for mycorrhizal races in a cheating orchid. Proc R Soc London Ser B Biol Sci 271:35–43. https://doi.org/10.1098/rspb.2003.2557

Taylor DL, Bruns TD, Leake JR, Read DJ (2002) Mycorrhizal specificity and function in myco-heterotrophic plants. In: Mycorrhizal ecology. Springer-Verlag, pp 375–413

Taylor DL, Bruns TD, Szaro TM, Hodges SA (2003) Divergence in mycorrhizal specialization within *Hexalectris spicata* (Orchidaceae), a nonphotosynthetic desert orchid. Am J Bot 90:1168–1179. https://doi.org/10.3732/ajb.90.8.1168

Taylor DL, McCormick MK (2008) Internal transcribed spacer primers and sequences for improved characterization of basidiomycetous orchid mycorrhizas. New Phytol 177:1020–1033. https://doi.org/10.1111/j.1469-8137.2007.02320.x

Tedersoo L, Bahram M, Zobel M (2020) How mycorrhizal associations drive plant population and community biology. Science 367:6480. https://doi.org/10.1126/science.aba1223

Tedersoo L, Brundrett MC (2017) Evolution of Ectomycorrhizal Symbiosis in Plants. In Biogeography of mycorrhizal symbiosis. Springer, Cham. pp. 407–467. https://doi.org/10.1007/978-3-319-56363-3_19

Tedersoo L, May TW, Smith ME (2010) Ectomycorrhizal lifestyle in fungi: global diversity, distribution, and evolution of phylogenetic lineages. Mycorrhiza 20:217–263. https://doi.org/10.1007/s00572-009-0274-x

Tedersoo L, Pärtel K, Jairus T, et al (2009) Ascomycetes associated with ectomycorrhizas: Molecular diversity and ecology with particular reference to the Helotiales. Environ Microbiol. https://doi.org/10.1111/j.1462-2920.2009.02020.x

Tedersoo L, Sánchez-Ramírez S, Kõljalg U, et al (2018) High-level classification of the Fungi and a tool for evolutionary ecological analyses. Fungal Divers 90:135–159. https://doi.org/10.1007/s13225-018-0401-0

Tedersoo L, Smith ME (2017) Ectomycorrhizal fungal lineages: detection of four new groups and notes on consistent recognition of ectomycorrhizal taxa in high-throughput sequencing studies. In: Biogeography of Mycorrhizal Symbiosis. pp 125–142. https://doi.org/10.1007/978-3-319-56363-3_6

Tedersoo L, Smith ME (2013) Lineages of ectomycorrhizal fungi revisited: Foraging strategies and novel lineages revealed by sequences from belowground. Fungal Biol Rev 27:83–99. https://doi.org/10.1016/j.fbr.2013.09.001

Tedersoo L, Suvi T, Beaver K, Saar I (2007) Ectomycorrhizas of *Coltricia* and *Coltriciella* (Hymenochaetales, Basidiomycota) on Caesalpiniaceae, Dipterocarpaceae and Myrtaceae in Seychelles. Mycol Prog. https://doi.org/10.1007/s11557-007-0530-4

Těšitelová T, Jersáková J, Roy M, et al (2013) Ploidy-specific symbiotic interactions: divergence of mycorrhizal fungi between cytotypes of the *Gymnadenia conopsea* group (Orchidaceae). New Phytol 199:1022–1033

Těšitelová T, Kotilínek M, Jersáková J, et al (2015) Two widespread green *Neottia* species (Orchidaceae) show mycorrhizal preference for Sebacinales in various habitats and ontogenetic stages. Mol Ecol 24:1122–1134. https://doi.org/10.1111/mec.13088

Těšitelová T, Tesitel J, Jersáková J, et al (2012) Symbiotic germination capability of four *Epipactis* species (Orchidaceae) is broader than expected from adult ecology. Am J Bot 99:1020–1032. https://doi.org/10.3732/ajb.1100503

Tondello A, Vendramin E, Villani M, et al (2012) Fungi associated with the southern Eurasian orchid *Spiranthes spiralis (L.) Chevall*. Fungal Biol 116:543–549. https://doi.org/10.1016/j.funbio.2012.02.004

Turenne CY, Sanche SE, Hoban DJ, et al (1999) Rapid identification of fungi by using the ITS2 genetic region and an automated fluorescent capillary electrophoresis system. Journal of Clinical Microbiology 37:1846–1851

Umata H (1995) Seed germination of *Galeola altissima*, an achlorophyilous orchid, with aphyllophorales fungi. Mycoscience 36:369–372. https://doi.org/10.1007/BF02268616

Umata H (1999) Germination and growth of *Erythrorchis ochobiensis* (Orchidaceae) accelerated by monokaryons and dikaryons of *Lenzites betulinus* and *Trametes hirsuta*. Mycoscience 40:367–371. https://doi.org/10.1007/BF02463883

Umata H (1997) Formation of endomycorrhizas by an achlorophyllous orchid, *Erythrorchis ochobiensis*, and *Auricularia polytricha*. Mycoscience 38:335–339. https://doi.org/10.1007/BF02464092

Umata H, Ota Y, Yamada M, et al (2013) Germination of the fully myco-heterotrophic orchid *Cyrtosia septentrionalis* is characterized by low fungal specificity and does not require direct seed-mycobiont contact. Mycoscience 54:343–352. https://doi.org/10.1016/j.myc.2012.12.003

UNITE Community (2019) UNITE USEARCH/UTAX release for Fungi. Version 18.11.2018. In: UNITE Community

van der Heijden MGA, Martin FM, Selosse MA, Sanders IR (2015) Mycorrhizal ecology and evolution: the past, the present, and the future. New Phytol 205:1406–1423. https://doi.org/10.1111/nph.13288

Vanegas-León ML, Sulzbacher MA, Rinaldi AC, et al (2019) Are Trechisporales ectomycorrhizal or non-mycorrhizal root endophytes? Mycol Prog 18:1231–1240. https://doi.org/10.1007/s11557-019-01519-w

Varga T, Krizsán K, Földi C, et al (2019) Megaphylogeny resolves global patterns of mushroom evolution. Nat Ecol Evol 3:668–678. https://doi.org/10.1038/s41559-019-0834-1

Veldre V, Abarenkov K, Bahram M, et al (2013) Evolution of nutritional modes of Ceratobasidiaceae (Cantharellales, Basidiomycota) as revealed from publicly available ITS sequences. Fungal Ecol 6:256–268. https://doi.org/10.1016/j.funeco.2013.03.004

Vujanovic V (2000) Viability Testing of Orchid Seed and the Promotion of Colouration and Germination. Ann Bot 86:79–86. https://doi.org/10.1006/anbo.2000.1162

Walker JF, Aldrich-Wolfe L, Riffel A, et al (2011) Diverse Helotiales associated with the roots of three species of Arctic Ericaceae provide no evidence for host specificity. New Phytol 191:515–27. https://doi.org/10.1111/j.1469-8137.2011.03703.x

Wang B, Qiu YL (2006) Phylogenetic distribution and evolution of mycorrhizas in land plants. Mycorrhiza 16:299–363. https://doi.org/10.1007/s00572-005-0033-6

Wang D, Jacquemyn H, Gomes SIF, et al (2021) Symbiont switching and trophic mode shifts in Orchidaceae. New Phytol 231:791–800. https://doi.org/10.1111/nph.17414

Wang Q, Guo L-D (2010) Ectomycorrhizal community composition of *Pinus tabulaeformis* assessed by ITS-RFLP and ITS sequences. Botany 88:590–595. https://doi.org/10.1139/B10-023

Wang X, Li Y, Song X, et al (2017) Influence of host tree species on isolation and communities of mycorrhizal and endophytic fungi from roots of a tropical epiphytic orchid, *Dendrobium sinense* (Orchidaceae). Mycorrhiza 27:709–718. https://doi.org/10.1007/s00572-017-0787-7

Warcup JH (1971) Specificity of mycorrhizal association in some australian terrestrial orchids. New Phytol 70:41–46. https://doi.org/10.1111/j.1469-8137.1971.tb02507.x

Waterman RJ, Bidartondo MI, Stofberg J, et al (2011) The effects of above-and belowground mutualisms on orchid speciation and coexistence. Am Nat 177:E54–E68. https://doi.org/10.1086/657955

Watkinson SC (2016) Mutualistic symbiosis between fungi and autotrophs. In: The Fungi. Elsevier, pp 205–243

Waud M, Busschaert P, Ruyters S, et al (2014) Impact of primer choice on characterization of orchid mycorrhizal communities using 454 pyrosequencing. Mol Ecol Resour 14:679–699. https://doi.org/10.1111/1755-0998.12229

Weiß M, Oberwinkler F (2001) Phylogenetic relationships in Auriculariales and related groups – hypotheses derived from nuclear ribosomal DNA sequences. Mycol Res 105:403–415. https://doi.org/10.1017/S095375620100363X

Weiß M, Selosse M-A, Rexer K-H, et al (2004) Sebacinales: a hitherto overlooked cosm of heterobasidiomycetes with a broad mycorrhizal potential. Mycol Res 108:1003–10. https://doi.org/10.1017/s0953756204000772

Weiß M, Waller F, Zuccaro A, Selosse M (2016) Sebacinales – one thousand and one interactions with land plants. New Phytol 211:20–40. https://doi.org/10.1111/nph.13977

White TJ, Bruns TD, Lee SB, et al (1990) Amplification and direct sequencing of fungal ribosomal RNA genes for phylogenetics. pp. 315–322 in Innis MA, Gelfand DH, Sninsky JS, and White TJ, eds. PCR protocols: a guide to methods and applications. Academic Press, New York

Whitridge H, Southworth D (2005) Mycorrhizal symbionts of the terrestrial orchid *Cypripedium fasciculatum*. Selbyana 328–334

Wijayawardene NN, Hyde KD, Rajeshkumar KC, et al (2017) Notes for genera: Ascomycota. Fungal Divers 86:1–594. https://doi.org/10.1007/s13225-017-0386-0

Wu G, Zhao H, Li C, et al (2015) Genus-wide comparative genomics of malassezia delineates its phylogeny, physiology, and niche adaptation on human skin. PLOS Genet 11:e1005614. https://doi.org/10.1371/journal.pgen.1005614

Xing X, Gao Y, Zhao Z, et al (2020) Similarity in mycorrhizal communities associating with two widespread terrestrial orchids decays with distance. J Biogeogr 47:421–433. https://doi.org/10.1111/jbi.13728

Xing X, Jacquemyn H, Gai X, et al (2019) The impact of life form on the architecture of orchid mycorrhizal networks in tropical forest. Oikos 128:1254–1264. https://doi.org/10.1111/oik.06363

Xing Y-M, Chen J, Cui J-L, et al (2011) Antimicrobial Activity and Biodiversity of Endophytic Fungi in *Dendrobium devonianum* and *Dendrobium thyrsiflorum* from Vietman. Curr Microbiol 62:1218–1224. https://doi.org/10.1007/s00284-010-9848-2

Xu J, Guo S (2000) Retrospect on the research of the cultivation of *Gastrodia elata* Bl, a rare traditional Chinese medicine. Chin Med J (Engl) 113:686–92

Xu JT, Guo SX (1989) Fungus associated with nutrition of seed germination of *Gastrodia elata* – *Mycena osmundicola* Lange. Acta Mycol Sin 8:221–226

Xu JT, Mu C (1990) The relation between growth of *Gastrodia elata* protocorms and fungi. Acta Bot Sin 32:26–31

Yagame T, Ogura-Tsujita Y, Kinoshita A, et al (2016) Fungal partner shifts during the evolution of mycoheterotrophy in *Neottia*. Am J Bot 103:1630–41. https://doi.org/10.3732/ajb.1600063

Yagame T, Yamato M (2013) Mycoheterotrophic growth of *Cephalanthera falcata* (Orchidaceae) in tripartite symbioses with Thelephoraceae fungi and *Quercus serrata* (Fagaceae) in pot culture condition. J Plant Res 126:215–222. https://doi.org/10.1007/s10265-012-0521-7

Yagame T, Yamato M, Mii M, et al (2007) Developmental processes of achlorophyllous orchid, *Epipogium roseum*: from seed germination to flowering under symbiotic cultivation with mycorrhizal fungus. J Plant Res 120:229–236. https://doi.org/10.1007/s10265-006-0044-1

Yamashita Y, Kinoshita A, Yagame T, et al (2020) *Physisporinus* is an important mycorrhizal partner for mycoheterotrophic plants: Identification of mycorrhizal fungi of three *Yoania* species. Mycoscience 61:219–225. https://doi.org/10.1016/j.myc.2020.05.003

Yeh CM, Chung KM, Liang CK, Tsai WC (2019) New insights into the symbiotic relationship between orchids and fungi. Appl Sci 9:1–14. https://doi.org/10.3390/app9030585

Yuan Z, Chen Y, Yang Y (2009) Diverse non-mycorrhizal fungal endophytes inhabiting an epiphytic, medicinal orchid *(Dendrobium nobile):* estimation and characterization. World J Microbiol Biotechnol 25:295–303. https://doi.org/10.1007/s11274-008-9893-1

Yukawa T, Ogura-Tsujita Y, Shefferson RP, Yokoyama J (2009) Mycorrhizal diversity in *Apostasia* (Orchidaceae) indicates the origin and evolution of orchid mycorrhiza. Am J Bot 96:1997–2009. https://doi.org/10.3732/ajb.0900101

Zelmer CD, Currah RS (1995) Evidence for a fungal liaison between *Corallorhiza trifida* (Orchidaceae) and *Pinus contorta* (Pinaceae). Can J Bot 73:862–866. https://doi.org/10.1139/b95-094

Zhang L, Chen J, Lv Y, et al (2012) *Mycena* sp., a mycorrhizal fungus of the orchid *Dendrobium officinale*. Mycol Prog 11:395–401. https://doi.org/10.1007/s11557-011-0754-1

Zhang W, Zhang X, Li K, et al (2018) Introgression and gene family contraction drive the evolution of lifestyle and host shifts of hypocrealean fungi. Mycology 9:176–188. https://doi.org/10.1080/21501203.2018.1478333

Zhao XL, Yang JZ, Liu S, et al (2014). The colonization patterns of different fungi on roots of Cymbidium hybridum plantlets and their respective inoculation effects on growth and nutrient uptake of orchid plantlets. World Journal of Microbiology and Biotechnology 30: 1993–2003. https://doi.org/10.1007/s11274-014-1623-2

Zhao R-L, Li G-J, Sánchez-Ramírez S, et al (2017) A six-gene phylogenetic overview of Basidiomycota and allied phyla with estimated divergence times of higher taxa and a phyloproteomics perspective. Fungal Divers 84:43–74. https://doi.org/10.1007/s13225-017-0381-5

Zhu GS, Yu ZN, Gui Y, Liu ZY (2008) A novel technique for isolating orchid mycorrhizal fungi. Fungal Divers 33:123–137

